# Fluorescence Lifetime Imaging in Plants: Practical guidelines for multiplexing, label-free imaging and data analysis

**DOI:** 10.64898/2026.06.11.731528

**Authors:** Beatrice Lace, Pascal Krohn, Morgane Batzenschlager, Vanessa Fieß, David Molina, Laura Ragni, Thomas Ott

## Abstract

Fluorescence Lifetime Imaging Microscopy (FLIM) is becoming a key technique for live-cell multiplexing and label-free detection of endogenous fluorescence in animal systems. Its potential in plant biology, however remains largely unexploited, despite its integration into a number of commercial microscopy setups. Here, we build a systematic, subcellular FLIM reference library for a panel of genetically-encoded fluorophores. Lifetime imaging of different fluorescent reporters targeted to distinct organelles (nucleus, plasma membrane, endoplasmic reticulum, etc.) and subsequent analysis of the decay curves using different modes allowed us to simultaneously discriminate up to four spectrally overlapping fluorophores solely by lifetime differences in specific subcellular compartments. Remarkably, fluorophores with lifetimes differing by as little as 0.1 ns can be reliably discriminated using one of these modes, namely Phasor-based analysis. Moreover, we show that the same fluorophores exhibit compartment-specific lifetime shifts, enabling Phasor separation of identical tags residing in different organelles. Finally, we extended the Phasor approach to label-free imaging of endogenous plant fluorescence. Together, these results establish FLIM-Phasor as a versatile, multiplex-capable tool for plant cell biology, opening new avenues for imaging strategies that yield higher content information at both cellular and tissue-level resolution.

## INTRODUCTION

Fluorescence imaging is a fundamental tool for investigating biological systems as it enables spatial and temporal resolution of dynamic processes in living cells and tissues. This is commonly achieved by labelling molecules, organelles, or cellular structures with fluorescent dyes or genetically encoded fluorescent proteins (FPs), allowing their localization and activity to be visualized with high sensitivity and minimal perturbation. The combination of multiple fluorescent tags further expands the analytical power by enabling the simultaneous observation of several molecular targets within the same sample. In conventional fluorescence microscopy, however, multiplexing typically relies on the spectral separation of fluorophores based on differences in their respective excitation and emission spectra (Lichtman and Conchello 2005). Therefore, spectral overlap restricts the number of reporters that can be reliably distinguished. In addition, endogenous cellular compounds often emit fluorescence within similar spectral ranges, complicating the unambiguous assignment of probe-derived signals (Lakowicz 2006). Fluorescence lifetime imaging microscopy (FLIM) overcomes these limitations by introducing fluorescence lifetime as an additional contrast dimension. The lifetime is the average time a fluorophore remains in the excited state before photon emission and represents an intrinsic property of each fluorescent species – its fingerprint (Berezin and Achilefu 2010). As such, it can be used to discriminate fluorophores with similar spectra and to separate probe-derived signals from endogenous fluorescence. In addition, lifetime signatures can also be exploited for label-free imaging of endogenous molecules, further reducing perturbation of biological systems (Datta et al. 2020). Moreover, because fluorescence lifetime is largely independent of fluorophore concentration, excitation intensity, and optical properties of the sample, it provides a robust parameter for quantitative analysis. Finally, its sensitivity to local environmental parameters such as viscosity, pH, solvent polarity and quenching makes it an excellent reporter of a fluorophore’s molecular surroundings (Berezin and Achilefu 2010).

Owing to these features and the increasing availability of commercial instrumentation, FLIM is rapidly emerging as a versatile imaging strategy across biological systems. Beyond its traditional use in Förster Resonance Energy Transfer (FRET) imaging (Wallrabe and Periasamy 2005), fluorescence lifetime has recently been exploited to extend the limits of live-cell multiplexing in animal systems. For instance, simultaneous imaging of nine genetically encodable FPs was successfully employed to track the entry of viral particles into human host cells, allowing to dissect the cellular remodelling occurring at different steps of this process (Starling et al. 2023). A similar resolution power in visualizing multiple subcellular targets was also achieved by using synthetic probes, either alone or in combination with engineered lifetime-modulating tags such as HaloTag or FAST (Frei et al. 2022a, 2022b; El Hajji et al. 2024). In parallel, FLIM is increasingly applied to assess cellular metabolism by resolving the endogenous fluorescence of coenzymes such as NADH and FAD (Lakowicz et al. 1992; Skala et al. 2007; Stringari et al. 2012). Such label-free approaches provide a real-time, non-invasive way to identify phenotypes in living cells and tissues without the application of dyes (Asadipour et al. 2025).

An important factor contributing to the increasing adoption of FLIM has been the introduction and further development of the Phasor approach for lifetime data analysis (Digman et al. 2008; Ranjit et al. 2018). Unlike conventional exponential fitting, which mathematically estimates decay parameters and requires an assumption on the number of species constituting a fluorescent signal, the Phasor method is fitting-free and does not require prior knowledge of the sample. Its beauty specifically lies in the conversion of the fluorescence decays into an intuitive 2D graphical representation, the Phasor plot. Following Fourier transformations, the pixels of an image are positioned in this semicircular plot according to the peculiar characteristics of their fluorescent signal, reflecting their molecular composition. Spatial regions with similar composition will cluster together, generating distinct and reproducible distributions, or clouds, whose position in the Phasor space serves as a molecular fingerprint. This representation enables, therefore, an immediate and intuitive assessment of fluorescence heterogeneity, allowing discrimination between molecular species and between mono- and multiexponential behaviours based solely on cluster position and shape. A key feature of this approach is the reciprocity between the FLIM image and the Phasor plot. Clusters in the plot can be selected and mapped onto the image to visualize their spatial distribution, while regions of the image can be analysed to determine their position in the Phasor space (Malacrida et al. 2021). Importantly, Phasor positions can be unambiguously described by a pair of coordinates. The Phasor plot acts, therefore, as an interactive map rapidly orienting the user across the complex raw lifetime data contained in an image while maintaining spatial resolution. These properties make the Phasor approach particularly powerful for precisely characterising fluorophore signatures and their shift upon changes in the molecular environment, and for resolving complex fluorescence signals, such as those arising from endogenous molecules (Malacrida 2023).

Despite its considerable potential, the application of FLIM and the Phasor approach in plant biology remains limited. This likely reflects both biological and technical constraints. The plant cell wall restricts the penetration of synthetic probes, limiting some experimental strategies (Colin et al. 2022), while FLIM itself requires specialized instrumentation and expertise, particularly for data analysis. In this regard, plant-focused guidelines exemplifying the use of lifetime imaging for multiplexing and endogenous fluorescence discrimination using the Phasor approach are currently lacking. Here, we aim at providing such practical insights, including a verified workflow to perform the downstream analyses using the Phasor approach. By systematically characterizing widely used genetically encoded FPs across three spectral channels, we demonstrate their combined use in live-cell multicolour FLIM in plants. Using reporters targeted to distinct subcellular compartments, we achieved simultaneous labelling of four cellular structures and show how lifetime differences can be resolved across a range of spatial relationships, from segregation to co-localization, using the Phasor approach. We also provide an example of how such concepts can be applied to resolve gene expression patterns using a multicolour promoter-reporter system in plant roots. We further define compartment-specific lifetime signatures and demonstrate that shifts as small as 0.1 ns are sufficient to distinguish the same fluorophore localized to different organelles - the nucleus and chloroplasts. In addition, we show how the complex endogenous fluorescence of plant cell walls, exemplified by the distinct lignin depositions across root tissues, can be characterized by its Phasor signature, representing a potential label-free strategy to identify phenotypes. Together, this work provides a practical framework for designing multicolour FLIM experiments in plants and for exploiting fluorescence lifetime as a molecular fingerprint to resolve both genetically encoded fluorescent reporters and endogenous signals in living plant cells and tissues.

## RESULTS

### Experimental strategy and downstream analysis to test multiplexing with genetically-encoded FPs in plant cells

To exemplify the use of FLIM to perform multiplexing in plant cells, we chose spectrally overlapping FP pairs in three different spectral regions (green, red and yellow), having either large or small differences in their lifetimes (Fig. 1A). These were targeted to specific cell compartments by fusing them to well-known organelle markers and transiently produced in *Nicotiana benthamiana* leaf epidermal cells (Fig. 1A). This system provided the required versatility to co-produce multiple combinations of fluorescent reporters with variable degrees of spatial overlap within single spectral channels (Fig. 1A). In order to systematically test the efficiency of lifetime-based separation of the different reporters, we followed a standard workflow for the acquisition and analysis of FLIM images (Fig. 1B). While imaging was performed using a Leica SP8 FALCON confocal platform, we used the offline LAS X Control Software version 4.8.2 for the subsequent analysis. This variant is part of the software suite developed for Leica’s new-generation STELLARIS microscopy platforms and includes an extended set of tools to analyse Phasor plots. For each pair of spectrally overlapping fluorescent reporters, we first show representative images of the intensity-only signal and of the FastFLIM (Fig. 1B, “Acquisition”). The latter is an imaging mode that generates fluorescence lifetime images in real time by directly measuring timing differences between excitation and detected fluorescence pulses, rather than reconstructing photon decay curves (Alvarez et al., 2019). These timing differences are converted into a lifetime value for each pixel (average photon arrival time) and then colour-coded according to a user-defined lifetime range to maximize contrast. At this stage, it does not require any mathematical fitting or Phasor analysis and thus allows the immediate assessment of lifetime differences while imaging the sample (Fig. 1B). To further analyse the lifetime data, we mainly adopted the Phasor approach, a fit-free method based on the Fourier transformation of the decay profile (extensively reviewed in Malacrida et al. 2021; Torrado et al. 2024). By mapping each pixel of the image on a semi-circular vectorial space according to its decay profile, the Phasor analysis provides a highly intuitive graphical representation of the composition and distribution of fluorescent species in the sample (a fingerprint, Fig. 1B). Pixels having similar decays (and composition of fluorescent species) will cluster together in an area defined by similar coordinates (G and S coordinates), forming clouds that can be reciprocally selected to colour-code the corresponding pixels in the image itself. This allows to promptly visualize the spatial distribution of the different fluorescent species within the image (overlay, Fig. 1B) and to separate their intensity contribution in sub-images, similarly to what mathematical fitting of the decay profile curve produces (separation, Fig. 1B). As a comparison, we provide the separation and lifetime quantification obtained via curve fitting in the supplementary figures. When using the Phasor approach and depending on the sample, we either analysed the fingerprint of the entire population of pixels (global) or of isolated region of interests (ROI) within the image (Fig. 1C, upper panel). We preferred the latter to determine the G and S coordinates of the fluorescent signal located in a specific region or compartment of the cell with the native function of the software, in order to avoid the influence of pixels with low photon counts (blue cloud, Fig. 1C, upper panel). In order to account for biological variability, we overlaid the Phasor fingerprints of all images from the same experimental series, or regions of interest within them, for each sample/cell analysed (Σ, Fig. 1C, upper panel). By doing this, we obtained a fingerprint describing the whole sample population, being it either heterogeneous (spread distribution) or homogeneous (confined cloud) (Fig. 1C, upper panels). According to this specific Phasor fingerprint, we then adopted different types of tools to colour code the images or separate the species (Fig. 1C, lower panels). Two fluorescent species having different lifetimes and being present at different locations within the cell (spatially separated) will originate discrete clouds at different positions in the Phasor plot. These can be selected with circular cursors to map the corresponding pixels in the image and separate the species (Fig. 1C, lower panel). When two fluorescent species spatially overlap within a region, the corresponding pixels will fall on the line (“mixing line”) connecting the Phasor position of the two pure species, at a location which reflects the fractional contribution of each species (Torrado et al. 2022). In this case, image segmentation can be obtained either by placing circular cursors on the Phasor position of the pure species, which further allows their separation into sub-images, or by using the “lifetime ruler” tool (Fig. 1C, lower panel). The latter is particularly useful when different blends of lifetimes are present within the sample population, resulting in a rather spread Phasor fingerprint where isolated clouds are not obviously distinguishable. Indeed, such an all-encompassing colour-coding allows a quick visualization of heterogeneous mixtures of two species and is useful to compare regions not only within a single image/cell but also across multiple images from the same sample/tissue, provided that the same ruler is applied to them (see Fig. 4 and Fig. 10). Importantly, the exact species composition of pixels clustering in a particular position along such extended fingerprint can be extracted using the ratio tool. This tool automatically computes the fractional contribution of the single species (Fig. 1C) and is particularly useful to assess the relative amount of two co-localizing fluorescent reporters (see Fig. 4 and Fig. S7).

**Figure 1.**
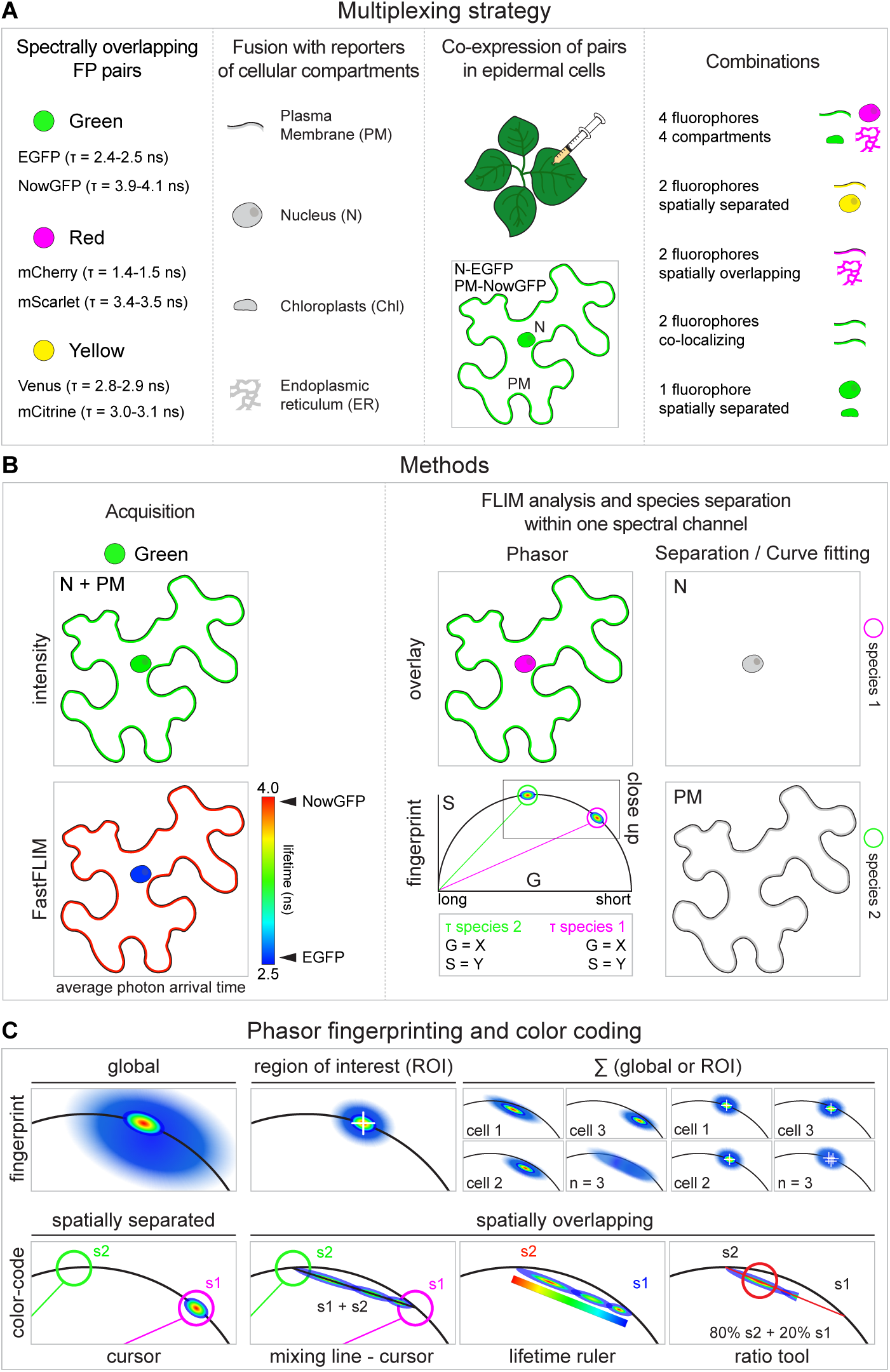
Graphical overview of the approach adopted to conduct multicolour lifetime imaging and Phasor analysis in plant cells. **A)** Green-, red- and yellow-emitting FP pairs with different lifetimes were targeted to different subcellular compartments by fusion with corresponding reporters and co-expressed in *N. benthamiana* leaf epidermal cells via agroinfiltration. Multiple combinations of spectrally overlapping fluorescent reporters were used to test lifetime-based separation across a range of spatial relationships. **B)** For each combination, we present representative images of the intensity-only signal and of the FastFLIM (the average photon arrival time), colour-coded according to a range of lifetimes appropriate to the FP pair (rainbow bar). FLIM images were further analysed with the Phasor approach, using the tools provided in the Leica LAS X Control Software (version 4.8.2). Fluorescent species were separated according to the position (defined by G and S coordinates and τ values) of the corresponding clusters in the Phasor fingerprint (often shown as close up). Their subcellular distribution pattern was visualized by overlaying the colour-code on the intensity signal (overlay) and/or by separating the intensity contribution of each species in sub-images. The results were compared to those obtained by curve fitting. **C)** For Phasor analysis, either the fingerprint of the whole image (global) or of a region of interest (ROI) was analysed. For ROIs, the mean G and S coordinates (white cross) of clouds were computed using the software. To obtain a representative Phasor fingerprint (Σ, n=3), we overlaid the individual fingerprints (cell 1, cell 2, cell 3) from the set of images (global), or ROI within them, from the same sample series. Different types of colour-codes were applied to the Phasor fingerprint depending on the spatial relationship between the two fluorescent species (s1 and s2) present in the sample. In case of spatial separation, each species will generate well-defined Phasor clouds which can be selected with circular cursors of different colours (green and magenta) to map the corresponding pixels in the image. In case of spatial overlap, pixels containing mixtures of the two species (s1+s2) will distribute on a straight “mixing line” (black line) joining the Phasor positions of the single species. Linear separation is obtained by placing the cursors outside the distribution, on or close to the semi-circle. Alternatively, the relative amount of the two species can be visualized by overlaying a lifetime ruler on the Phasor distribution or quantified using the ratio tool, which computes the ratio of the two species at a chosen position (red circular cursor) along the mixing line (red line). See main text and Materials and Methods for more details.

### Simultaneous discrimination of four fluorescent reporters combining spectral resolution and fluorescence lifetime information

Among the genetically-encoded FP variants that have been developed over the years, EGFP and mCherry are the most widely adopted ones. Due to the limited overlap of their excitation and emission spectra, they are often used in combination to design dual-colour reporters. In order to generate a more complex four-colour reporter, we additionally selected NowGFP and mScarlet as fluorophores to combine with EGFP and mCherry, respectively, owing to their overlapping spectral properties but different lifetimes (EGFP = 2.4-2.5 ns; NowGFP = 3.9-4.1 ns; mCherry = 1.4-1.5 ns; mScarlet 3.4-3.5 ns, (Seefeldt et al. 2008; George Abraham et al. 2015; Sarkisyan et al. 2015; Bindels et al. 2017, 2020; Lace et al. 2023). Since the capacity to resolve two fluorescent reporters within the same spectral channel using FLIM relies on a homogeneous fluorescence lifetime distribution, we first measured the lifetimes of all four FPs when being targeted to different intracellular compartments, the nucleus (N) and the plasma membrane (PM). This was achieved by fusing them to either a histone protein (H3.1 or H3.3) from *Medicago truncatula* or a remorin protein (REM1.3) from *Arabidopsis thaliana*, respectively (Fig. S1A-D). An additional factor to consider in FLIM imaging is the repetition rate of the pulsed laser. This parameter determines the time between the pulses (12.5 ns for 80 MHz, 25 ns for 40 MHz, 50 ns for 20 MHz). To avoid pulse pile-up, leading to artificially shorter measured lifetimes, the repetition rate should be longer than the fluorescence decay time of the fluorophore (greater than five times the lifetime of a fluorophore). Here, we conducted measurements using 20, 40 or 80 MHz as repetition rates (Fig. S1A-D). Considering the lifetimes of the chosen FPs, we expected 20 MHz to be sufficient to map the entire intensity decay of all four FPs, 40 MHz to be suitable for EGFP, mCherry and mScarlet and 80 MHz for EGFP and mCherry. Interestingly, however, we found no statistically significant differences between the lifetime determinations at 20 and 40 MHz for all four FPs, including NowGFP, in both compartments. Instead, all green- and red-emitters, including the short-lived mCherry, showed statistically significant shorter lifetimes when imaged using a 80 MHz laser repetition rate (Fig. S1A-D). Therefore, we decided to use 40 MHz to conduct all the following lifetime imaging experiments with these genetically-encoded FPs as probes. In addition, we noticed that all FPs exhibited shorter and more homogeneous lifetimes when targeted to the PM compared to the nucleus (Fig. S1A-D). This suggests that the peculiar characteristics of these two compartments, and/or the behaviour of the fusion protein within them, might influence the FP’s lifetime. Despite this variability, the lifetime differences between green- and red-emitting pairs were still in the range being sufficient to allow their discrimination (Δτ EGFP/NowGFP ≈ 1.5 ns; Δτ mCherry/mScarlet ≈ 2.0 ns, Fig. 2A). In order to test this, we imaged *N. benthamiana* leaf epidermal cells simultaneously producing the four FPs targeted to four different subcellular compartments (PM, chloroplasts, nucleus and endoplasmic reticulum, Fig. 2A). Indeed, we could efficiently discriminate the two FP pairs according to the average photon arrival time recorded by FastFLIM, with the two subcellular compartments within the same spectral channel being colour-coded differently (Fig. 2B). To further segment the image, we applied the Phasor approach (Fig. 2C-D). First, we colour-coded the intensity signal of each channel via overlay according to its Phasor fingerprint. The green-emitting FPs showed two main discrete and distant clouds, whose pixels could be clearly mapped to the chloroplasts (Chl, EGFP, green, τ = 2.203 ns) and the PM (NowGFP, cyan, τ = 3.955 ns) within the corresponding image (Fig. 2C-D). The regions of close apposition between the two subcellular compartments, where the signal of the two FPs co-occurs, could additionally be identified by selecting the distributions mapping on the mixing line connecting the main Phasor clusters, at positions reflecting the predominance of NowGFP (purple, τ = 3.352 ns) or EGFP (orange, τ = 2.533 ns) (Fig. 2D). For the red-emitting fluorophores, only the ER-localized mCherry defined a discrete Phasor cloud (yellow, τ = 1.372 ns, Fig. 2C-D), while the nuclear mScarlet rather mapped at different positions along the mixing line connected to the mCherry cloud (magenta and red, τ = 3.002 ns and 2.475, Fig. 2C-D). This is consistent with the different degrees of spatial overlap of the ER and the nucleus. To further resolve the spatial distribution of the four different fluorescent species, we separated each of them in a sub-image according to the cursors on the Phasor (Fig. 2C and 2E). This resulted in the efficient and unambiguous isolation of each subcellular compartment (PM, chl, N, ER) targeted by one of the four fluorescent reporters. Finally, we performed image segmentation by curve fitting, applying either a bi- or a triexponential model to fit the global decay (Fig. S2A-B). Using the two-components fit, neither the subcellular distribution patterns nor the lifetime values obtained for each fluorescent reporter matched the expectations (Fig. S1A-D and Fig. S2A). This discrepancy was solved by increasing the number of fitting components to three. However, this “created” a third image for each channel, with lifetimes and spatial distributions that could not be unambiguously assigned to any of the reporters (Fig. S2B). Based on these data, we conclude that NowGFP/EGFP and mCherry/mScarlet are suitable FP pairs when being used in combination with Phasor analysis to precisely and unambiguously discriminate four genetically-encoded fluorescent reporters directed to different subcellular targets within the same cell.

**Figure 2.**
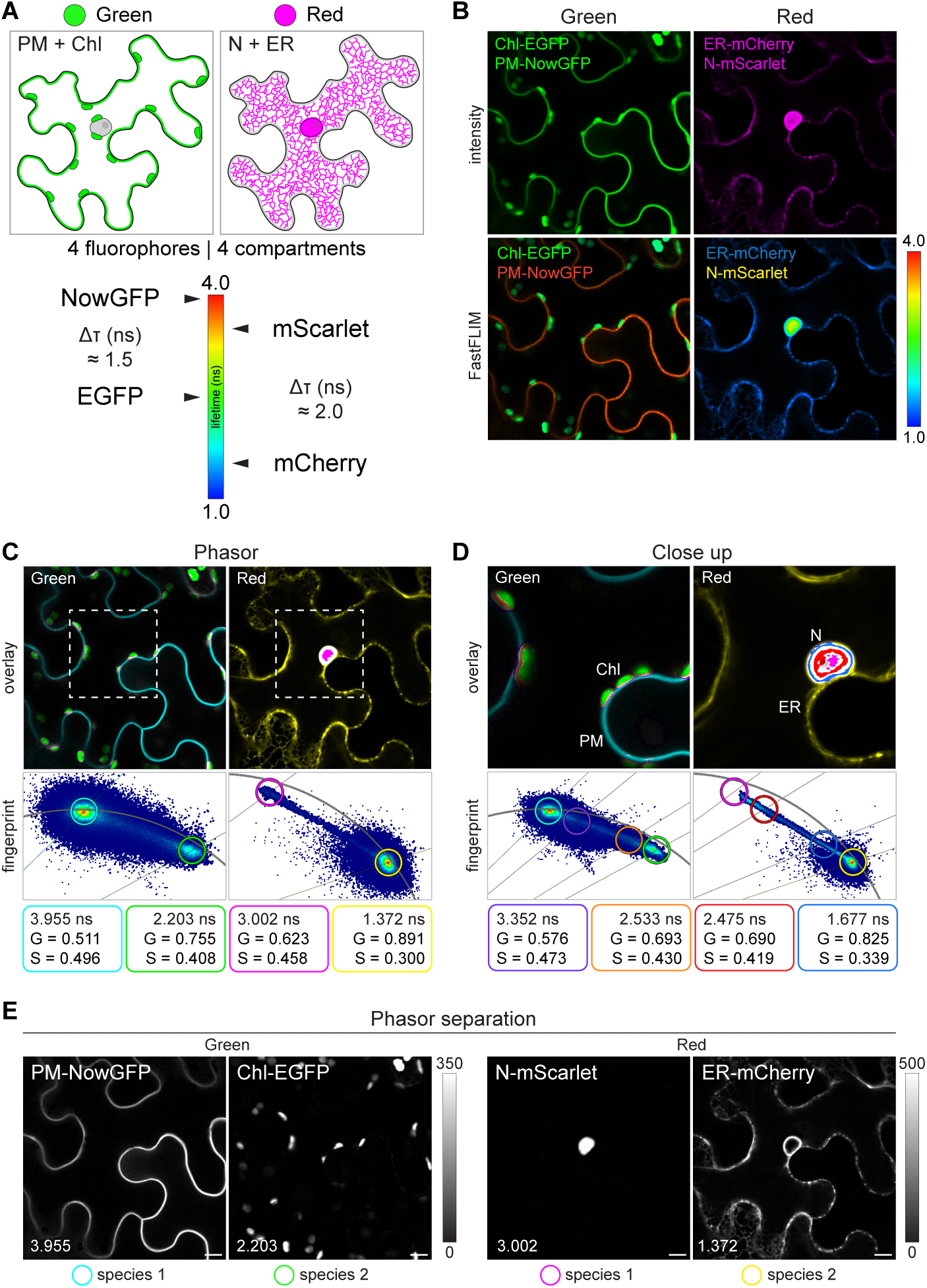
Live-cell multicolour lifetime imaging with two pairs of fluorescent reporters in different spectral windows. **A)** Two green- (EGFP and NowGFP) and two red- (mCherry and mScarlet) emitting FPs with large lifetime differences (Δτ) were targeted to four different compartments of *N. benthamiana* leaf epidermal cells (schematic). PM = plasma membrane; Chl = chloroplasts; N = nucleus; ER = endoplasmic reticulum. **B)** Intensity-only and FastFLIM confocal images of an epidermal cell showing the discrimination of two fluorescent reporters within the green and red channel based on lifetime values (rainbow bar). **C)** Identification of the fluorescent reporters and mapping of their subcellular distribution based on the Phasor fingerprint. The white dashed box indicates the region shown in the close up in (**D**). **D)** Regions of spatial overlap between the reporters are shown in the same cell imaged at higher magnification. **E)** Separation of the intensity contribution from each fluorescent reporter in individual sub-images according to the Phasor cursors in (**C**). Images of the isolated species are shown in gray with the minimum and maximum intensity value for the green and the red channel indicated by the corresponding calibration bar. The G and S coordinates together with the lifetime values (ns) of the cursors used to analyse the Phasor fingerprints in (**C**) and (**D**) are provided. Images are representative of two independent replicates with at least 6 cells analysed. Scale bar = 10 μm.

### Phasor separation of spectrally overlapping fluorescent reporters having a high degree of spatial overlap or similar lifetimes

Next, we aimed to test whether a higher degree of spatial co-occurrence of two reporters would also allow their unambiguous discrimination. For this, we combined the ER-localized mCherry (SP-AtWAK2-mCherry-HDEL) with a PM-localized mScarlet (mScarlet-AtREM1.3) and imaged agroinfiltrated *N. benthamiana* epidermal cells constitutively producing them (Fig. 3A). In cells co-producing the two reporters, the average arrival time recorded at the ER was shifted from the expected lifetime values of mCherry due to the co-occurrence with the PM-associated mScarlet, but the two compartments were still readily distinguishable (Fig. 3B). Such a shift was evident when comparing the colour-code of the ER in a neighbouring cell, where the signal of the PM reporter was not detectable (Fig. 3B). To perform image segmentation via Phasor, we overlaid the fingerprints of six different images exhibiting a heterogenous combination and spatial overlap of the two reporters in the epidermal cells (Ʃ fingerprints, Fig. 3C). This not only allows to display the variability within the samples, as all pixels from every image are mapped on the Phasor plot, but also to define the Phasor positions of the two fluorescent components present in the system without having an “a priori” knowledge. This is obtained by placing Phasor cursors at the positions where the mixing line intersects the semicircle (magenta and green cursors, Fig. 3C) and is useful in order to apply the same segmentation to every image belonging to the sample/condition. By using this method, we could efficiently separate the ER-localized mCherry from the PM-associated mScarlet (Fig. 3C). This demonstrates that Phasor-FLIM is suitable to resolve two co-occurring reporters tagged with FPs having distant lifetimes such as the mCherry/mScarlet pair. Fitting the decay profile with either a bi- or a triexponential model also allowed us to discriminate the two fluorescent species and assign them to the expected subcellular compartment, but yielded either imprecise lifetime values or a third short-lived fluorescent species, similar to what we previously observed (Fig. S3A-B and S2A-B).

**Figure 3.**
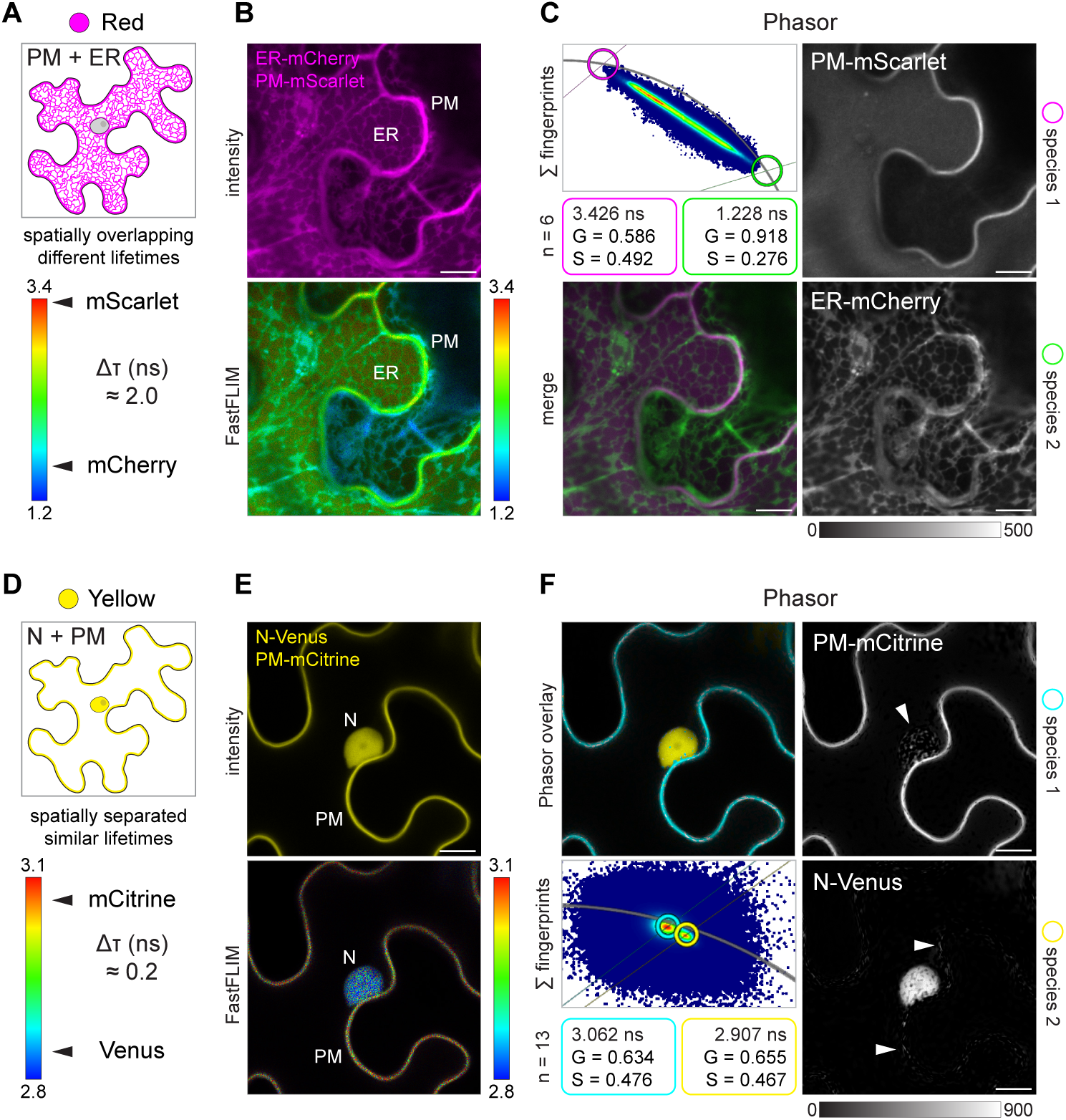
Phasor separation of spatially overlapping fluorescent reporters or with small lifetime differences. **A)** Two red-emitting FPs (mCherry and mScarlet) with a large lifetime difference (Δτ) were targeted to two spatially overlapping compartments of *N. benthamiana* leaf epidermal cells (schematic). PM = plasma membrane. ER = endoplasmic reticulum. **B)** Intensity-only and FastFLIM confocal images of epidermal cells showing the discrimination of the two fluorescent reporters based on lifetime values (rainbow bar). **C)** Phasor separation of the two reporters based on the overlaid fingerprints from multiple images (Σ fingerprint). The positions of the single species (green and magenta circles) were obtained by placing the corresponding cursors outside the distribution, at the intersection between the mixing line and the Phasor semicircle. Images of the isolated species are shown in gray with the minimum and maximum intensity value indicated by the calibration bar. The image showing the merge is also provided. **D)** Two yellow-emitting (mCitrine and Venus) FPs with a small lifetime difference (Δτ) were targeted to two spatially separated compartments of *N. benthamiana* leaf epidermal cells (schematic). N = nucleus; PM = plasma membrane. **E)** Intensity-only and FastFLIM confocal images of an epidermal cell showing the discrimination of the two fluorescent reporters based on lifetime values (rainbow bar) **F)** Phasor separation of the two reporters mapping to close but consistently distinct positions in the Phasor plot across cells (Σ fingerprints). Images of the isolated species are shown in gray with the minimum and maximum intensity value indicated by the calibration bar. White arrowheads indicate misassigned pixels. The G and S coordinates together with the lifetime values (ns) of the cursors used to analyse the Phasor fingerprint are provided. n = number of images. Data are from two independent replicates. Scale bar = 10 μm.

The next question we wanted to address is whether FPs having closer lifetime values could also be used for multiplexing. For this, and in order to cover an additional window of the visible spectrum, we selected the yellow-emitters Venus (τ = 2.8-2.9 ns, Denay et al. 2019) and mCitrine (τ = 3.0-3.1 ns, Söhnel et al. 2016; Lace et al. 2023), which are both widely adopted EYFP-derived variants with enhanced brightness (Fig. 3D, https://www.fpbase.org/). To test their separation, we targeted them to the nucleus or the PM via fusion with a nuclear localization signal (NLS-3xVenus) or with the high-affinity phosphatidylinositol 4-phosphate biosensor 2xPH^FAPP1^ (Simon et al. 2014) (mCitrine-2xPH^FAPP1^), respectively (Fig. 3D). First, we characterized the decay of the two fluorescent reporters when being expressed singularly in epidermal cells using Phasor analysis (Fig. S4A). Both of them showed a homogenous and stable decay across samples, as shown by the corresponding overlay of Phasor fingerprints, with mean lifetime values of 2.862 ns for the nuclear-localized 3xVenus (yellow, Fig. S4A) and of 3.083 ns for the PM-localized mCitrine (cyan, Fig. S4A), confirming the small difference in between their lifetimes (Δτ = 0.2 ns, Fig. 3D). In addition, we observed a slight shift of the Venus cluster towards the inside of the semicircle, which is consistent with the multiexponential decay previously reported for this fluorophore (Fig. S4A, Sarkar et al. 2009). When co-expressed in the same cells, the two reporters could be discriminated by FastFLIM. However, the colour-coding of each of the two targeted compartments resulted non homogeneous due to the narrow lifetime range (Fig. 3E). Despite these close lifetime values, the two fluorescent reporters appeared as two distinct clouds in the Phasor plot, enabling to clearly map and distinguish the nuclear-localized triple Venus (yellow, τ = 2.907 ns) from the PM-localized mCitrine reporter (cyan, τ = 3.062 ns) on the image (Fig. 3F). This was further confirmed by analysing the Phasor fingerprints of region of interests (ROIs) selected on the nucleus (ROI1) or on the PM (ROI2) within the same set of images (Fig. S4B). The two species could also be successfully isolated in separate images, although some pixels of the nucleus labelled by Venus were assigned to the image of the PM-Citrine, and vice versa (Fig. 3F). While the segmentation via Phasor worked successfully, fitting of the decay curve with a biexponential model did not separate the signal of the two fluorophores sufficiently (Fig. S4C). This indicates that the fit-free Phasor approach is more suited to discriminate FPs with small differences in lifetime values.

### Phasor discrimination and quantitative estimation of co-localizing fluorescent reporters within one spectral channel

While visualizing the co-localization of two fluorescent reporters in the same cellular compartment and/or tissue is a key application in plant cell biology, the estimation of the relative amount of two spatially overlapping proteins is essential for the use of fluorescent biosensors assessing, for instance, cell cycle stages or cell polarity (Simon et al. 2014; Tejos et al. 2014; Otero et al. 2016; Doumane et al. 2021). Typically, the use of spectrally distinguishable FPs (e. g. EGFP/mCherry, CFP/YFP) strictly limits the number of channels available for combinations with other probes. In order to overcome this limitation, we exploited the difference in fluorescence lifetime of EGFP and NowGFP (Fig. 2A and 4A) as an example to test whether spectrally overlapping FP pairs enable multiplexing of co-localizing markers or reporters. Thus, we co-expressed two membrane-associated fusion proteins (NowGFP-REM1.3 and EGFP-SYMREM1, Marín et al. 2012; Jarsch et al. 2014) in *N. benthamiana* leaves (Fig. 4A). In order to assess whether the lifetime information could be leveraged to determine the differential distribution of the two PM-associated reporters in a tissue, we imaged the boundaries between multiple epidermal cells exhibiting different degrees of expression of the two reporters and selected suitable cells by the FastFLIM colour-coding (Fig. 4B-C). We then acquired pairs of images by focusing on the cell surface (tangent plane, Fig. 4B-C) and at a plane crossing the cell volume (secant plane, Fig. 4B-C). Cells producing either EGFP-SYMREM1 only (cell1, Fig. 4B) or NowGFP-REM1.3 only (cell2, Fig. 4B), or various combinations of them (cell1, cell3 and cell4, Fig. 4C), could be readily distinguished according to the colour-code of their individual PMs visualized at the tangent plane. The lifetime recorded at the border between the cells, where the PMs lay in close proximity, corresponded to a combination of the lifetimes of the individual cells (arrowheads, Fig. 4B-C). To confirm this and further characterize the differential PM labelling of cells, we overlaid the Phasor fingerprints of the entire set of images (Fig 4D and Fig. S5A). This resulted in pixels clustering at different regions along the mixing line connecting the NowGFP-REM1.3 only (red, τ = 3.992 ns) and the EGFP-SYMREM1 only (blue, τ = 2.414 ns) positions, reflecting the different proportions of the two fluorescent reporters present across all the images (Fig. 4D). By placing a lifetime ruler along the mixing line, we defined an all-encompassing colour-coding to apply to the Phasor fingerprint of any of the images, enabling the prompt visualization of the composition and spatial distribution of the two reporters (Fig 4D-E and 4G). Phasor analysis further allowed to attribute and map onto the images their relative PM enrichment (Fig. 4F and 4H), as well as to visualize it by separating each reporter and its intensity contribution into individual images according to the Phasor cursors (Fig 4I-L and Fig. S5B). Altogether, we show that the relative amount of two genetically-encoded fluorescent reporters targeted to the same subcellular compartment can be rapidly and efficiently visualized and analysed in cells and tissues using the Phasor approach using a single spectral channel. Such an approach not only frees other spectral channels for the monitoring of additional probes, but also offers a convenient alternative to quantify the extent of co-localization of two fluorescent reporters at cellular resolution.

**Figure 4.**
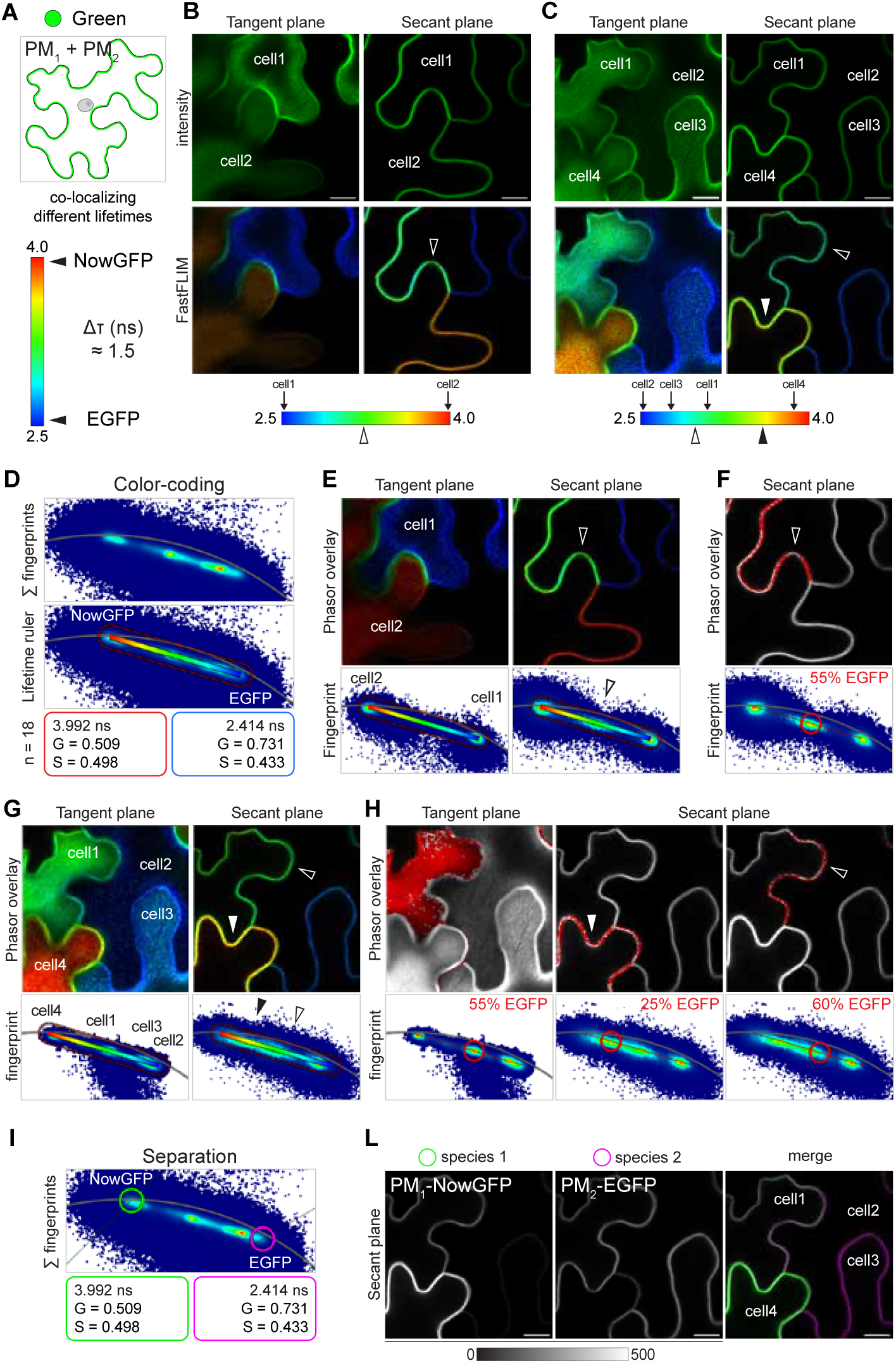
Phasor separation and quantitative estimation of co-localizing fluorescent reporters. **A)** Two green-emitting FPs (EGFP and NowGFP) with a large lifetime difference (Δτ) were targeted to the same compartment of *N. benthamiana* leaf epidermal cells. PM = plasma membrane. **B-C)** Intensity-only and FastFLIM confocal images of the boundaries between multiple epidermal cells acquired at tangent and secant planes. Cells (cell1-2 in (**B**) and cell1-4 in (**C**)) and borders between them (empty and solid arrowheads) are mapped on the lifetime rainbow bar according to the colour shown in the FastFLIM image. **D)** Overlay of the Phasor fingerprints from all the images of the sample series (Σ fingerprints). An all-encompassing colour-code was defined by positioning a lifetime ruler along the mixing line joining the NowGFP and EGFP positions. **E and G)** Evaluation and comparison of the PM-resident FP composition of each cell (tangent plane) and of the border between them (secant plane, empty and solid arrowheads) by overlaying the same lifetime ruler to the Phasor of single images. **F and H)** Mapping and quantification of the relative intensity contribution (expressed as percentage, %) of each reporter at different PM regions using the ratio tool. **I-L)** Separation of the intensity contribution from each fluorescent reporter in individual sub-images by placing circle cursors at the NowGFP and EGFP positions. Images of the isolated species are shown in gray with the minimum and maximum intensity value indicated by the calibration bar. The G and S coordinates together with the lifetime values (ns) of the lifetime ruler and cursors used to analyse the Phasor fingerprint are provided. n = number of images. Data are from two independent replicates. Scale bar = 10 μm.

### A multicolour transcriptional reporter of plant host infection by bacteria

To test the applicability of multicolour FLIM to a more complex biological system, we designed a three-colour transcriptional reporter to co-visualize the expression pattern of two genes during symbiotic infection of *M. truncatula* roots by fluorescent bacteria at cellular resolution (Fig. 5A). In such a bipartite system, it is crucial to simultaneously co-monitor multiple host cellular targets and the bacterial symbionts in order to study the cellular mechanisms enabling the accommodation of microbes within the plant cells and tissues. To exemplify the method, we chose the infection-specific genes *RPG* and *VPY,* which are required to steer bacterial colonization across plant host tissues (Arrighi et al. 2008; Murray et al. 2011). This process is initiated in curled root hairs (RH, Fig. 5A) where a host-derived tubular structure, the infection thread (IT, Fig 5A), forms and progresses intracellularly across cortical cell layers (cortex, Fig 5A), while being colonized by dividing bacteria. We generated transcriptional reporters of the two genes by driving tandem EGFP or mScarlet to the nucleus using a Nuclear Localization Signal (NLS), respectively (proMtRPG::NLS-2xEGFP in blue and proMtVPY::NLS-2xmScarlet in red, Fig. 5A). These were combined with a constitutively expressed ER-localized NowGFP (proAtUbi::SP-AtWAK2-NowGFP-HDEL, green, Fig 5A) serving both to track actively growing IT structures (Fournier et al. 2008) and to select positively transformed roots. Composite plants (i.e. plants with a WT shoot and a transgenic root system) were then inoculated with a compatible bacterial strain constitutively expressing mRFP1 (*Ensifer medicae* mRFP1, light blue, Fig. 5A) and imaged at 4-6 days post-inoculation. Live-cell imaging using the FastFLIM mode allowed to readily distinguish the two spectrally overlapping fluorescent reporters within each channel according to their lifetime differences (Fig. 5B). This revealed a strong and specific co-activation of the *RPG* and *VPY* promoters in cells on the so-called IT trajectory (Batzenschlager et al. 2025), the path an actively growing IT takes through root cortical cells, clearly indicated by the cytoplasmic ER accumulation around the tubular structure colonized by fluorescent bacteria (Fig. 5B, white arrowheads). A shorter lifetime was associated to the ER-NowGFP in these infected cells compared to the surrounding ones, likely due to spatial overlap with the NLS-2xEGFP reporter accumulating in the cytoplasm as a result of high activity of the *RPG* promoter (Fig. 5B and Fig. S6A, green channel). Consistently, signal corresponding to the NLS-2xmScarlet reporting on the VPY promoter activity was also visible in the cytoplasm of the first cortical cell crossed by the IT (Fig. 5B, red channel). Since the two transcriptional reporters emit in separate channels, their individual nuclear signals and spatial overlap were also easy to visualize using standard intensity-based confocal imaging (Fig. 5C). In this case, however, they could not be separated from the signal of the ER and the bacteria, respectively. Precise isolation of each reporter in individual images was instead possible by analysing the lifetime data using the Phasor approach (Fig. S6A and Fig. 5D). This confirmed the strong co-expression of the two genes on the IT trajectory marked by the ER and by the bacteria, consistent with the function of their encoded products in mediating IT polar growth (Liu et al. 2019; Lace et al. 2023). Since the nuclei of two neighbouring cells were also labelled by the transcriptional reporters (Fig. 5D, asterisks), we performed FLIM imaging at deeper focal planes to check for the possible presence of other infection events (Fig. S6B). Indeed, we found a second IT penetrating the tissues ahead of these cells, further supporting the IT-associated transcriptional co-activation of *RPG* and *VPY* (Fig. S6B). Since IT polar growth also requires a third genetic component called *LIN* (Kuppusamy et al. 2004; Li et al. 2023), we decided to generate a four-colour reporter containing an additional cassette for the expression of a nuclear-localized tandem mCherry under the control of the LIN promoter (proMtLIN::NLS-2xmCherry). Such a design would enable us to further test the separation of co-localizing fluorescent reporters and the simultaneous resolution of five targets. Co-activation of the *LIN* and *VPY* promoters in cells on the IT trajectory should indeed result in shortening of the lifetime measured in the corresponding nuclei compared to the three-colour reporter. As expected, standard confocal imaging of infection events at 4-6 dpi with *E. medicae* mRFP1 did not allow to resolve such co-activation, due to spectral overlap of the mCherry and mScarlet transcriptional reporters (Fig. S7A). Instead, this was clearly shown by the colour-code of nuclei in the FastFLIM image (Fig. S7B) and by the analysis of the Phasor fingerprint (Fig. S7C), indicating a higher expression of *VPY* over *LIN* in the cell contacted by the IT compared to the neighbouring one (Fig. S7B-C, N1 and N2 respectively). Phasor separation further allowed to compare the expression patterns of the three co-expressed genes, suggesting a strong co-upregulation of *VPY* and *RPG* in cell preparing to host bacterial symbionts (Fig. S7D, N1). While the signal of the mRFP1-tagged bacteria could not be isolated in an individual image, it could clearly be separated from the mCherry and mScarlet signal in the Phasor overlay (Fig. S7C). This confirms the previously reported possibility to discriminate three overlapping spectral FPs when spatially separated (Aoyama et al. 2023) and further exemplifies how live-cell multicolour imaging of five different targets can be achieved. Altogether, our results demonstrate that multicolour FLIM based on genetically encoded fluorescent reporters can be successfully applied to investigate biological processes occurring in plants at cellular resolution.

**Figure 5.**
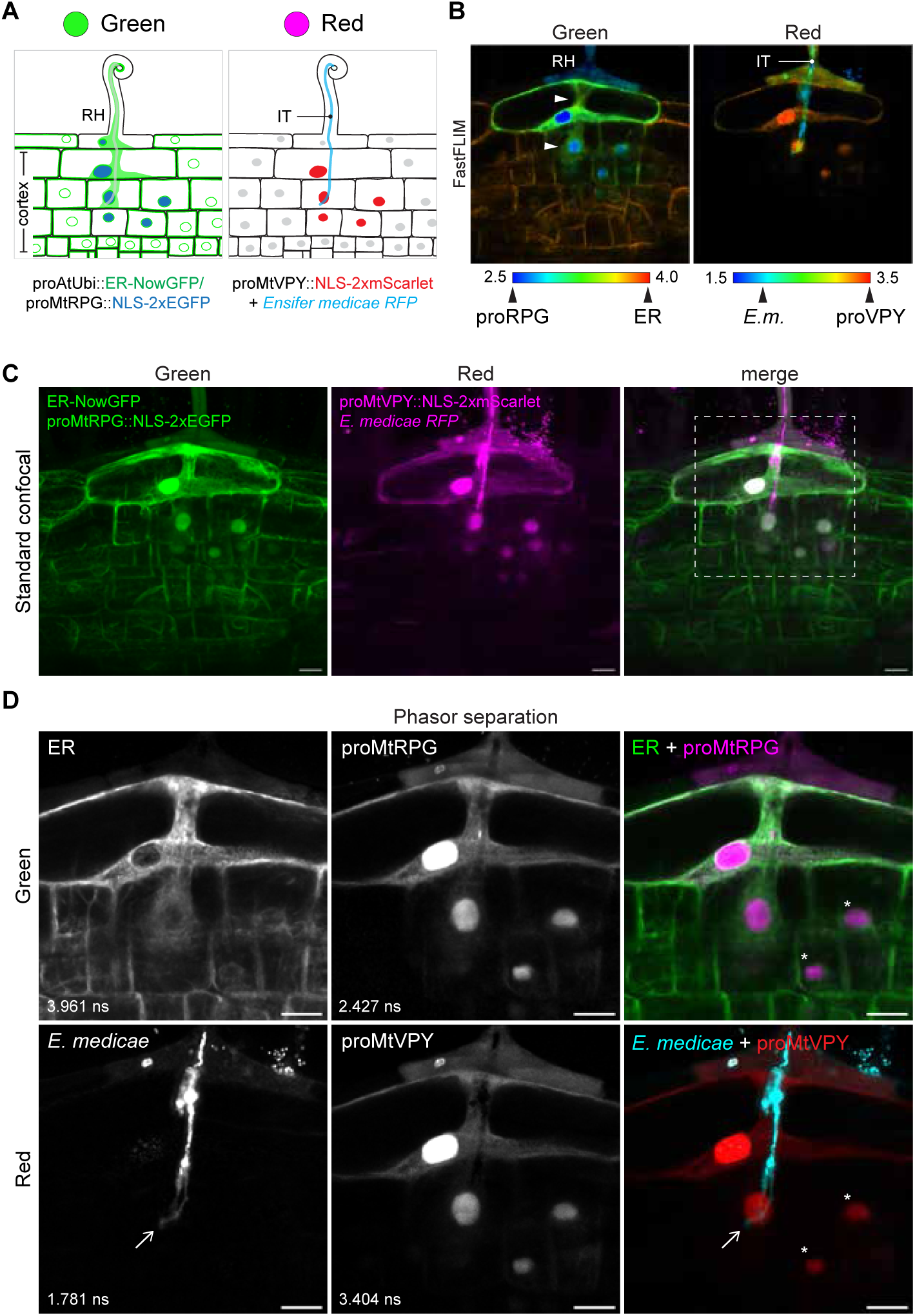
A three-colour transcriptional reporter enabling the tracking of gene expression patterns during bacterial colonization of *M. truncatula* roots. **A)** Graphical sketch representing the different targets labelled by the multicolour transcriptional reporter during bacterial infection in the green and red channel. RH = root hair, IT = infection thread. **B)** Live-cell FastFLIM confocal image showing the co-activation of *RPG* (proRPG) and *VPY* (proVPY) promoters in cells crossed by an actively growing IT colonized by fluorescent bacteria and surrounded by ER accumulations (white arrowheads). The four fluorescent reporters are mapped on the lifetime rainbow bar according to their expected lifetimes. *E.m*. = *Ensifer medicae* C) Maximum intensity projections of a z-stack (37 steps, step size = 1 μm) obtained by imaging at a lower magnification the same infection event as in **(A)** using standard confocal microscopy. The white dashed box in the merge depicts the area which was subsequently imaged using FLIM shown in (**D**) and in Fig. S6A. **D)** Separation of the intensity contribution from each fluorescent reporter in individual sub-images according to the Phasor cursors in Fig. S6A. Arrows indicate the tip of the IT structure colonized by bacteria. Asterisks indicate nuclei located in cells adjacent to the IT trajectory. Images of the isolated species are shown in gray. A minimum and a maximum intensity value of 0 and 800 or 200 were used to optimize the brightness and contrast of the green and the red channel, respectively. Merges are colour-coded according to the Phasor cursors in Fig. S6A. Lifetime values (ns) of the isolated species are annotated on the corresponding images. Images are representative of one independent replica with at least 5 infection events analysed. Scale bar = 20 μm.

### Compartment-specific lifetime signatures of green- and red-emitting fluorophores

Our initial measurements indicated that a shift occurs to an FP’s lifetime depending on its subcellular localization (Fig. S1A). In order to evaluate this more comprehensively, we used the Phasor approach to characterize and compare the decay of EGFP when targeted to different subcellular compartments (nucleus, ER, PM, mitochondria, peroxisomes, chloroplasts) or when freely diffusing in the cytoplasm and nucleus of *N. benthamiana* epidermal cells (Fig. 6A and Fig. S8). By overlaying the fingerprints of ROIs selected on a set of images acquired for every EGFP-tagged organelle marker (Σ fingerprints, Fig. 6A) and additionally plotting those on a graph (Prism, Fig. 6A), we determined the mean positions of the corresponding clusters, together with their lifetime values. Such analysis showed compartment-specific lifetime signatures for this widely adopted green emitting FP (Fig. 6A-C). Indeed, lifetimes values ranged from an average of 2.592 ns when EGFP was targeted to the nucleus to an average value of 2.140 ns when the fluorophore localized to chloroplasts, and a maximum lifetime difference (maxΔτ) of ≈ 0.4 ns (Fig. 6A-C). While the EGFP decay mapped to similar positions of the Phasor plot when recorded in most of the compartments (Fig. 6B-C), which indeed appeared colour-coded similarly in the FastFLIM images (Fig. S8), it was consistently shifted towards shorter lifetimes when analysed in chloroplasts, as also reflected by the FastFLIM colour-coding (Fig. S8). Moreover, the cloud of the chloroplasts was shifted towards the inside of the plot, indicating a multiexponential decay of the FP in these photosynthetic organelles. This is conceivable given the complex and highly active metabolic environment of these organelles, differing significantly from the cytoplasm (Jarvis and López-Juez 2013). We then asked whether such a shift is specific to EGFP or if it is exhibited by other green- or red-emitting FPs. For this, we applied the same approach to analyse the decay of NowGFP and mCherry targeted to either the nucleus, the PM or the chloroplasts (Fig. 7A-D and S9A-B). For both FPs, we observed a shift of the Phasor cloud representing the decay at ROIs selected on the chloroplasts compared to those selected on the nucleus and the PM, with the nuclear-localized reporters exhibiting the longest lifetime (Fig. 7A-D and S9A-B), as observed for the EGFP (Fig. 6A). Interestingly, the long-lived NowGFP exhibited the most pronounced lifetime difference (maxΔτ ≈ 0.8 ns) compared to the medium-lived EGFP (maxΔτ ≈ 0.4 ns) and the short-lived mCherry (maxΔτ ≈ 0.1 ns). This chloroplasts-specific lifetime shortening could be due to quenching of the FPs due to energy transfer to the chlorophyll, to the spatial overlap with the endogenous fluorescence of such plant-specific molecule, or to the microenvironmental parameters characterizing the thylakoid membranes or stroma (viscosity, refractive index, pH; Tregidgo et al. 2008; Shen et al. 2013; Tamada et al. 2014; Kirchhoff 2019). To investigate the possible underlying reasons, we first further dissected the decay of the three chloroplast-localized fluorescent reporters by comparing the Phasor fingerprints of ROIs selected on stromules, which are stroma-filled membrane-bound projections extending from the chloroplast surface, to the ones obtained at the organelle body (Fig. S10A-C). We found that all FPs exhibited a longer decay in the stromules (orange, Fig. S10A-C) compared to the body (green, magenta and cyan, Fig. S10A-C) where thylakoid membranes are located. The lifetime difference was more pronounced in case of the green-emitters EGFP (Δτ ≈ 0.2 ns) and NowGFP (Δτ ≈ 0.4 ns) compared to the red-emitter mCherry (Δτ ≈ 0.06 ns) (Fig. S10A-D). Although longer than those measured in the body, the lifetimes measured in the stromules were still shorter compared to those calculated for the three FPs when localized in other compartments (Fig. 6A-C and 7A-D). This suggests that indeed not only the peculiar environment associated to the thylakoid-rich body but also the characteristics of the stroma may influence the decay of FPs, especially of the green-emitters. Lastly, to consider the potential influence of the endogenous fluorescence, we analysed the Phasor fingerprint of chloroplasts in *N. benthamiana* untransformed cells imaged using the same settings as used for the green- and red-emitting fluorophores (Fig. S11). The endogenous signal exhibited a specific and very rapid decay in both the green (τ = 0.122 ns) and the red (τ = 0.367 ns) channel (Fig. S11), with values in the range of those previously reported in other systems (Singhal and Rabinowitch 1969; Krause and Weis 1991; Berezin and Achilefu 2010). However, when analysing the overlay of Phasor fingerprints, we found that the cloud of the fluorescently-labelled chloroplasts did not precisely lay on the mixing line connecting the positions of the endogenous fluorescence and of the stromules, suggesting that its shift cannot be explained, or at least not fully, by the spatial overlap with the endogenous fluorescence of the organelles (Fig. S11), but it may rather be due to a combination of different factors. Importantly, the short-decaying endogenous fluorescence of the chloroplasts, although fainter, could be efficiently discriminated and isolated from the signal of a freely-diffusing EGFP located in both the nucleus and the cytoplasm of transiently transformed *N. benthamiana* epidermal cells, by both FastFLIM and Phasor analysis (Fig. S12A-C). Finally, Phasor analysis also allowed to identify and separate the decay of the same FP when targeted to the chloroplasts and to the nucleus of the same epidermal cells (Fig. 8A-C and S13A-F). This was achieved by the specific signature of the FPs when localized to these two different subcellular compartments. Altogether, we show that the lifetimes of both green- and red-emitters can vary depending on their subcellular localization, providing a quantitative estimation of such variability and therefore establishing a useful reference for imaging genetically-encoded fluorescent reporters in plant cells. Our observations are not only coherent with the sensitivity of fluorophores decay to the molecular environment but they are also relevant, for instance, in the context of quantitative studies such as FLIM-FRET, relying on a precise estimation of the donor lifetime.

**Figure 6.**
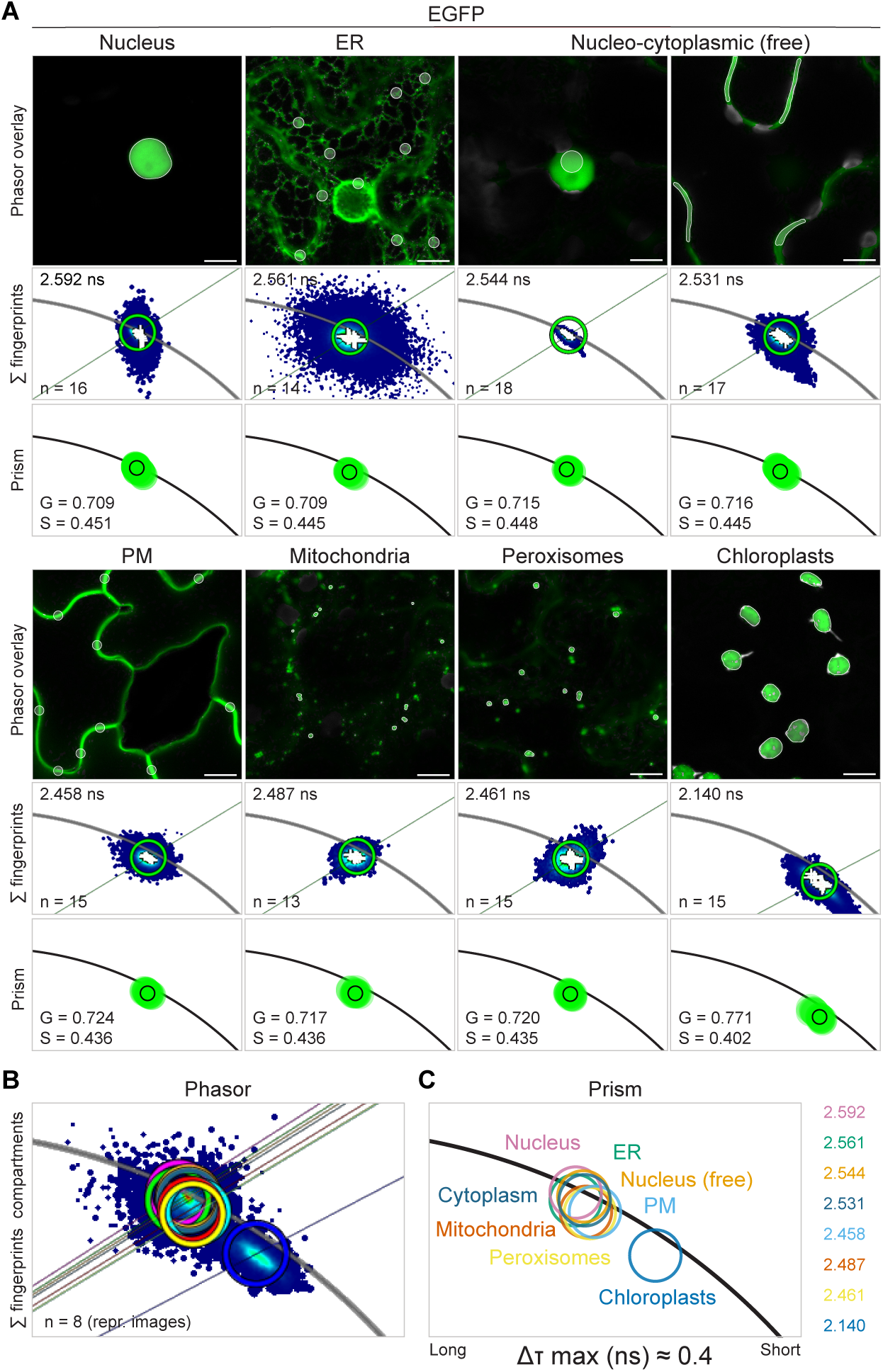
Lifetime signatures of EGFP in different subcellular compartments. **A)** Phasor analysis and representative confocal images of EGFP targeted to different subcellular compartments of *N. benthamiana* leaf epidermal cells. For each reporter, ROIs (white outlines) were selected on the labelled compartment to obtain its Phasor fingerprint. The Phasor fingerprints of ROIs from the whole set of images were overlaid (Σ fingerprints) and a green cursor was centered on the resulting cluster to obtain its lifetime value (ns) and to colour-code the representative image. The mean G and S coordinates (white crosses) of each ROI retrieved from the LAS X SP8 Control Software were additionally plotted on a graph (green solid dots) using GraphPad Prism. The smaller green dot outlined in black represents the average G and S calculated in Excel, whose values are reported on the graph. **B-C)** The Phasor positions of EGFP in the different compartments (circles of different colours) are shown together by overlaying the Phasor fingerprints of the representative images (**B**) or by plotting the average G and S on a graph (**C**). The lifetime values of the single compartments and the largest lifetime difference (Δτ max) between them are reported. n = number of images. Data are from at least two independent replicates. Scale bar = 10 μm.

**Figure 7.**
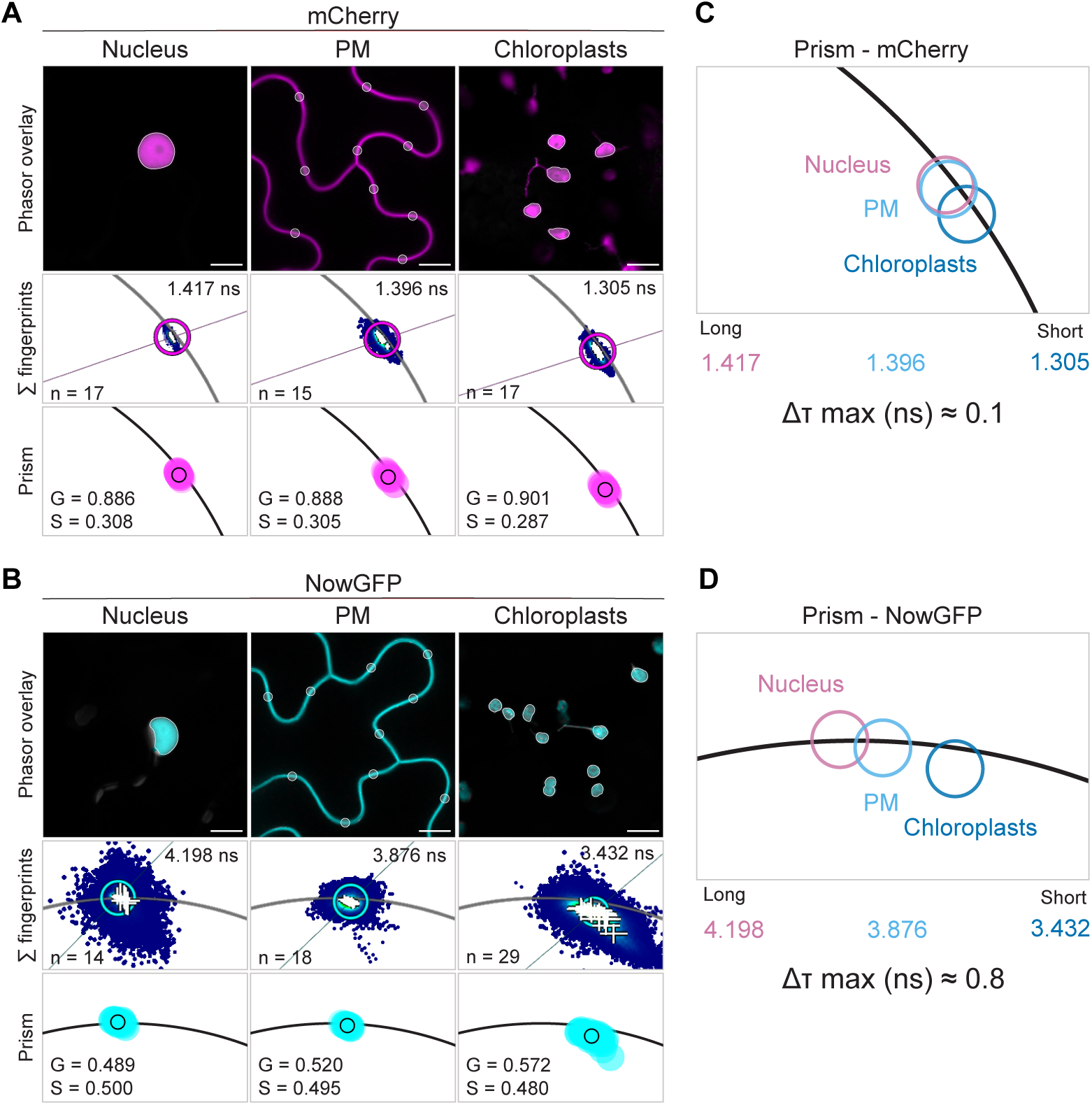
Chloroplast-specific lifetime shifts of NowGFP and mCherry. **A-B)** Phasor analysis and representative confocal images of mCherry (magenta) and NowGFP (cyan) targeted to the nucleus, PM and chloroplasts of *N. benthamiana* leaf epidermal cells. The analysis was carried out as for EGFP (see legend Figure 5). **C-D)** Graphs showing the average G and S of mCherry and NowGFP in the different compartments. The lifetime values (ns) of the single compartments and the largest lifetime difference (Δτ max) between them are reported. Data are from at least two independent replicates. Scale bar = 10 μm.

**Figure 8.**
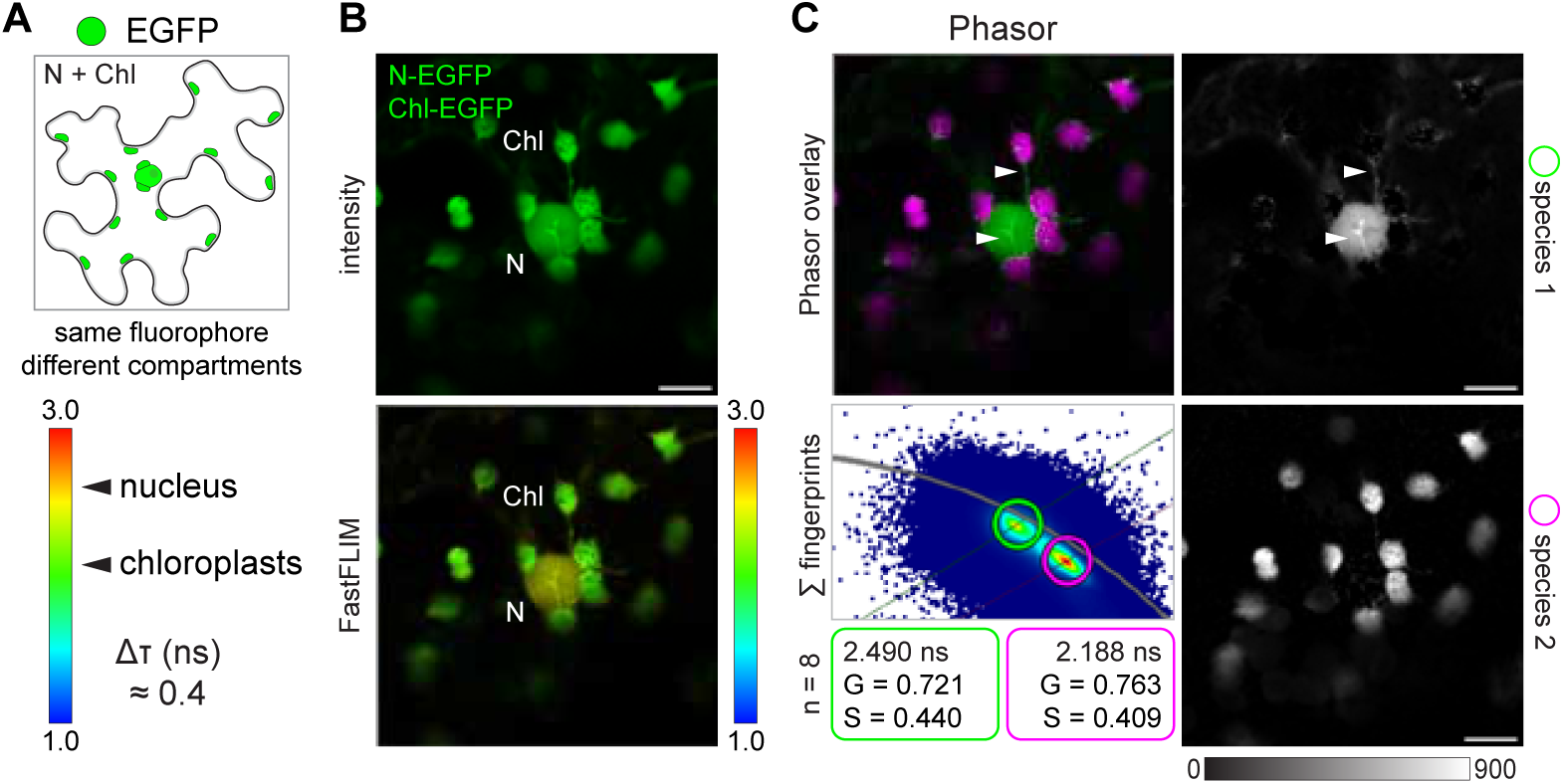
Phasor separation of the same FP targeted to different subcellular compartments. **A)** EGFP was simultaneously targeted to the nucleus (N) and the chloroplasts (Chl) of *N. benthamiana* leaf epidermal cells to test if the lifetime shift (Δτ) supports discrimination of the two compartments. **B)** Intensity-only and FastFLIM confocal images of an epidermal cell showing the discrimination of the same FP based on lifetime values (rainbow bar). **C)** Phasor separation of the chloroplasts- and nuclear-localized EGFP mapping to close but consistently distinct positions in the Phasor plot across cells (Σ fingerprints). Arrowheads point at stromules, whose signal segregates together with the nuclear-localized EGFP. Images of the isolated species are shown in gray with the minimum and maximum intensity value indicated by the calibration bar. The G and S coordinates together with the lifetime values (τ) of the cursors used to analyse the Phasor fingerprint are provided. n = number of cells. Data are from two independent replicates. Scale bar = 10 μm.

### Label-free characterization of the endogenous fluorescence (autofluorescence)

Besides the chlorophyll contained in chloroplasts, plants accumulate many other “autofluorescent” compounds, among which the aromatic cell wall polymer lignin is one of the most abundant. While its endogenous fluorescence can be problematic and confounding when overlapping with the emission of a fluorescent reporter, it can also serve for the label-free detection of its spatial patterning and composition in tissues using specific imaging techniques. Indeed, lifetime imaging has been proposed as a method to characterize the composition of lignin in wood cell walls (Donaldson and Radotic 2013; Chabbert et al. 2018; Wightman et al. 2019). However, its use to define the depositions occurring in roots has only been partially explored (Escamez et al. 2021). Since such characterization could represent a useful read-out in both developmental and genetic studies, we analysed lignin as an example of an autofluorescent structure in plant cells. First, we asked whether specific decays can be associated with the different lignin impregnations occurring in root tissues and whether we can describe a reference “Phasor fingerprint” to use for comparative analysis. For this, we imaged cross-sections of tomato (*Solanum lycopersicum*) roots obtained from the region where lignin can be detected in the exodermis, endodermis and xylem (Manzano et al. 2024, Fig. 9A). The tomato root is a particularly suitable model for studying lignin depositions, as the endodermis is not suberized and suberin deposition in the exodermis occurs only in the most mature regions of the root. Thus, it provides access to numerous cells that are exclusively lignified, avoiding confusion between the autofluorescence of the two polymers (Cantó-Pastor et al. 2024; Manzano et al. 2024). Since lignin has a broad fluorescence emission range and can be excited with both UV and visible light (Donaldson 2020), we decided to use a 488 nm laser line for excitation and a 500-550 nm emission window, which would allow to simultaneously image, and possibly discriminate, an EGFP-tagged reporter using the same channel. Using these spectral parameters and imaging the root tissues at different magnifications, we were able to clearly visualize the typical patterns of lignified root cell walls, i.e. the tracheary elements of the primary xylem in the central cylinder, the Casparian strips forming the apoplastic barrier of the endodermis, and the U-shaped polar lignin cap (PLC) on the epidermal face of exodermal cells (Fig. 9A-B). The presence of lignin at these specific sites was further confirmed by basic fuchsin staining combined with the counterstaining of cellulosic cell walls with calcofluor white, carried out on the same sections after imaging the endogenous fluorescence (Fig. 9B, right column). Although differences were noticeable in the intensity-only images, with a generally stronger signal associated to the Casparian strips and the corner regions within the xylem when compared to the exodermis, they did not allow *per se* to discriminate between the lignin depositions occurring at the different cell wall domains (Fig. 9B, left column). Instead, specific decays were promptly revealed by the FastFLIM colour-coding. Here, a shorter and homogenous lifetime was associated with the endodermal Casparian strips compared to other lignified cell walls (Fig. 9B, middle column). In addition, clear lifetime differences were exhibited by the central six-cell files of the xylem compared to the terminal two to three cell files, possibly reflecting the developmental transition between meta- and protoxylem, respectively (Fig. 9B, middle column). By performing three-dimensional (3D) reconstruction of Z-stacks from the central cylinder and the entire cross-section, we further confirmed the spatial distribution of these specific decays. We could recognize the typical ring-like structure of the Casparian strips, with polymerized lignin deposited on the radial and transverse walls of the root endodermis, as well as the peculiar shapes of the xylem vessels and of the exodermal barrier (Fig. 9C-D). As a next step, we generated a deeper characterization and description of the lignin patterns using the Phasor approach. For this, we first obtained a representative Phasor fingerprint by overlaying the Phasors from the images of multiple cross sections (depicting all the tissues) from different plants (Fig. 10A). The pixels distributed along a curved line located inside the right part of the Phasor plot, indicative of complex multiexponential decays (Fig. 10A). To describe such blend of decays, we placed a lifetime ruler along the pixel cloud, identifying a range of lifetimes spanning from 0.237 ns to 1.107 ns, with intermediate values placed at positions where apparent pixel clusters were recognizable (Fig. 10A). Applying such colour-code to single sections, allowed a fast visualization of similarities and differences across the samples. We found a uniform short-lived decay at the Casparian strips domain, a long-lived decay associated to the metaxylem compared to a shorter one characterising the protoxylem, while a rather variable lifetime patterning was present at the exodermal PLC (Fig. 10B). To confirm and further characterise these tissue-specific decays, we analysed the Phasor fingerprints of isolated regions within the cross-sections imaged at higher magnification (Casparian strips, xylem, central cylinder + endodermis, exodermis, Fig. 10C). This revealed a rather confined cluster for the endogenous signal of the Casparian strips (τ = 0.422 ns, cyan, Fig. 10C left panel), well distinguishable from the more spread fingerprint of the xylem vessels (τ = 0.585 – 1.107 ns, red to greenish cyan, Fig. 10C middle panels), suggesting differences in composition/organization between the two distinct lignin depositions. The decay of the exodermal PLC mainly distributed at intermediate positions (τ = 0.585 – 0.745 ns, greenish cyan to green, Fig. 10C right panel, plant 1 and plant 8), with slightly longer decays associated to a broader lignification of the anticlinal cell walls (yellow or red, Fig. 10C right panel, plant 8), possibly reflecting different developmental stages of this peripheral barrier. By mapping the Phasor position of ROIs selected on the proto- and metaxylem, we could further define the differences between the two vascular tissues, with the protoxylem decay consistently clustering at shorter lifetimes (τ = 0.821 ns, cyan, Fig. 10D) compared to the decay of the metaxylem (τ = 0.927 ns, yellow, Fig. 10D). Altogether, we demonstrate that the lignin depositions occurring at different cell wall domains of tomato roots are associated to specific decays and provide a method to efficiently identify, characterize and discriminate their endogenous fluorescence using Phasor analysis. Label-free approaches, as exemplified for lignin in tomato roots here, open promising avenues to describe, characterize and compare the changes associated with lignified cell walls following genetic or chemical manipulation.

**Figure 9.**
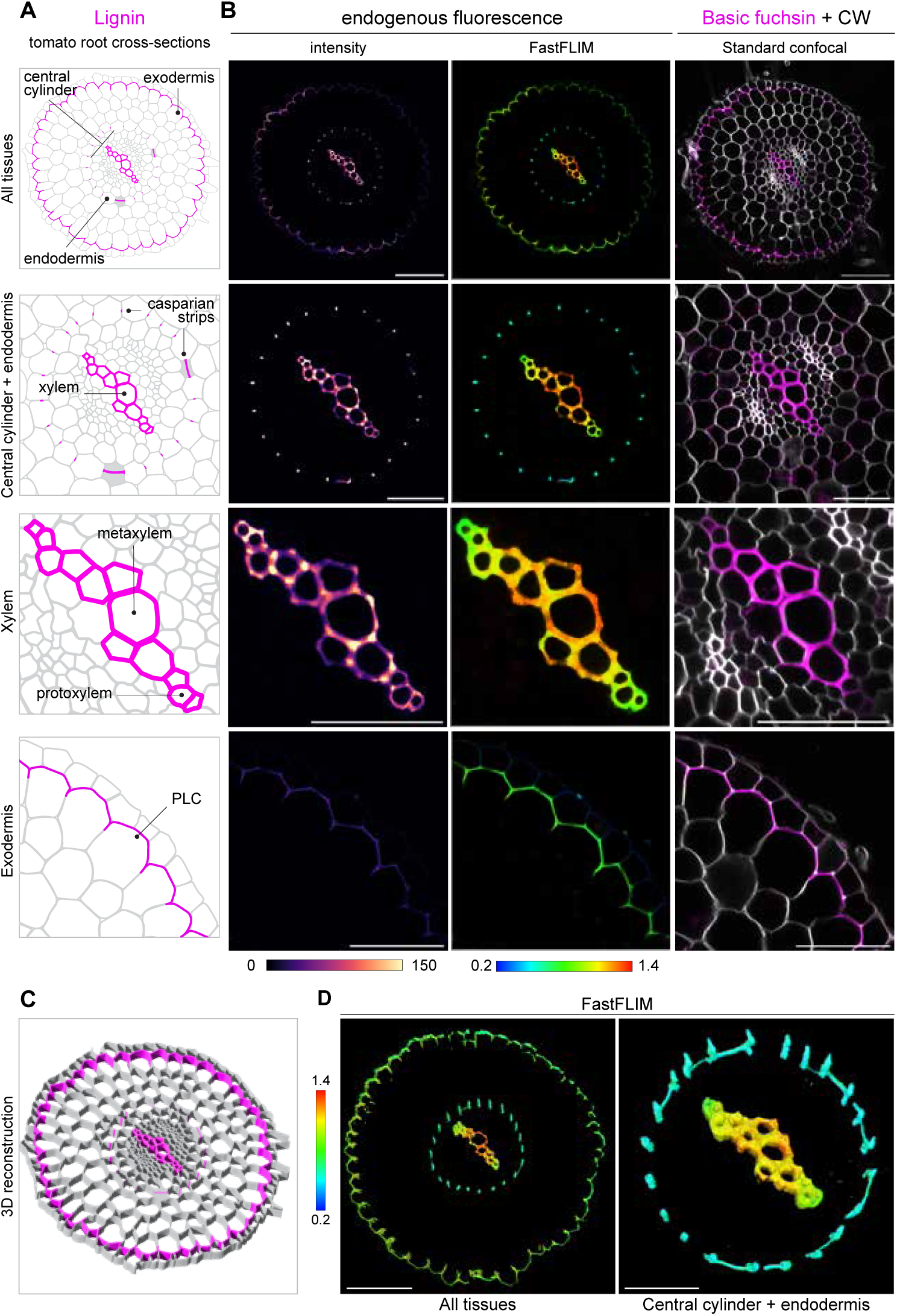
Specific fluorescence lifetimes of lignin depositions revealed by FLIM in tomato root tissues. **A)** Graphical representations of lignin impregnations (magenta) occurring at the cell wall of different tissues (indicated on the sketches) in tomato root cross-sections. PLC = polar lignin cap. **B**) Confocal images of different regions of a cross-section showing the endogenous fluorescence of tissues as intensity-only signal and FastFLIM, alongside the basic fuchsin (magenta)/calcofluor white (CW, gray) dual staining of lignin and cellulose. Intensity and FastFLIM images are colour-coded with a magma or rainbow look-up table, respectively, with the minimum and maximum intensity or lifetime values indicated by the corresponding calibration bars. **C-D**) 3D reconstructions visualizing the spatial distribution of lignified cell walls in cross-sections of tomato roots as a graphical representation (**C**) or as FastFLIM confocal images of the endogenous fluorescence (**D**). Images in (**D**) are of two different cross-sections. Images are representative of two independent biological replicates. Scale bar = 100 μm (all tissues), 50 μm (central cylinder + endodermis, xylem, exodermis).

## DISCUSSION

The availability of commercial instruments with integrated FLIM technology, together with the introduction of the Phasor approach to analyse lifetime data, is leading to the rapid adoption of FLIM as a fundamental imaging tool. Among its features, the capacity to discriminate spectrally overlapping fluorophores based on their lifetime signatures offers the possibility to co-visualize a higher number of subcellular targets compared to standard fluorescence imaging, and therefore probe biological mechanisms with an enhanced level of information and contrast. To facilitate the adoption of such technology by the plant community, where its use is still limited, we present here practical guidelines and examples on how to perform live-cell multicolour lifetime imaging at cellular and subcellular resolution, using the Phasor approach to analyse lifetime images in a systematic and reproducible manner. We show how three genetically encoded FP pairs in different spectral channels can be efficiently separated based on their Phasor fingerprint (Fig. 2-4) and demonstrate that the combination of green- and red-emitting pairs enables the simultaneous visualization of up to four cellular targets in a plant cell (Fig. 2). We further prove that these concepts can be successfully applied to perform live-cell multicolour FLIM not only in *N. benthamiana* leaves under ectopic expression conditions but also in complex biological systems such as the intracellular colonization of legume roots by symbiotic bacteria (Fig. 5). By designing multicolour transcriptional reporters, we exemplify the use of FLIM to monitor gene expression patterns coinciding with ongoing cell remodelling processes, achieving resolution of five different cellular targets (Fig. 5 and Fig. S7). Yellow-emitters such as Venus and mCitrine are often used as tags in plant imaging due to the limited autofluorescence of plant tissues in this range (Donaldson 2020). We show that Phasor analysis can resolve them thanks to their narrow lifetime distributions and despite the small difference in their lifetime (Fig. 3D-F). This pair could potentially be combined with blue- and red- emitting pairs to further increase the multiplexing degree up to six targets. Due to the limited excitation range of our confocal FLIM platform, with 470nm being the lowest available wavelength, we could not directly test blue-emitters. mTFP1 and mTurquoise2, however, were shown to be a suitable pair in animal systems (Starling et al. 2023), with a reported lifetime difference (1.3 ns) in a similar range of the FP pairs we tested here. While the Venus-mCitrine pair is best suited to label spatially separated structures due to its low dynamic range, we demonstrate that FPs exhibiting larger lifetime differences as the mScarlet/mCherry and EGFP/NowGFP pair are effective for multicolour strategies that require the monitoring of co-occurring or co-localizing molecular targets (Fig. 3A-C and Fig. 4A-L). By mapping multiple images on the Phasor plot, we present how the Phasor positions of the individual fluorescent reporters can be easily identified, enabling a reproducible and effective image segmentation, together with the quantification of the relative contribution from each reporter at regions where they overlap (Fig. 3C and Fig. 4D-L). This enables the generation of single-colour markers or biosensors, in analogy to the cell-cycle indicators based on red-emitting FPs (FUCCI-Red, Shirmanova et al. 2021) or HaloTag variants (LT-Fucci, Frei et al. 2022a) designed for mammalian systems. As an alternative application, we demonstrate the possibility to visualize and quantify with cellular resolution the differential but overlapping expression patterns of two genes using one spectral channel and labelling one cellular compartment (Fig. S7). In general, for the design of multicolour tools it is important to consider that lifetime-based separation of fluorescent probes within a single spectral channel depends on the relative number of photons collected from each species (Frei et al. 2022a; Rahim et al. 2022). This, in turn, is determined by both their intrinsic brightness and their abundance in the sample, as their signals are acquired simultaneously using a single excitation wavelength. In practice, differences in brightness - such as those reported for the mCherry/mScarlet pair (https://www.fpbase.org/compare/mcherry,mscarlet/) - can be compensated by appropriate experimental design. For example, the dimmer fluorophore (mCherry) can be used to label the more abundant target, while the brighter fluorophore (mScarlet) can be assigned to less abundant structures (Fig. 2A-E, ER and nucleus). Similarly, the relative abundance is influenced, and can be modulated, by the strength of the promoters driving the expression of the reporter genes. On the other hand, targeting two transcriptional reporters to the same compartment represents a way to directly compare gene expression levels (Fig. S7). Importantly, the quantification of the fractional contributions using the Phasor approach is intensity-dependent, as it reflects the relative photon contribution of each fluorophore at a given pixel (Torrado et al. 2022). Therefore, for co-localization approaches, it is advisable to use fluorophores with similar brightness—such as EGFP and NowGFP (https://www.fpbase.org/compare/egfp,nowgfp/)—so that variations of the Phasor position more directly reflect differences in abundance rather than intrinsic photophysical properties. To counterbalance the different brightnesses of mCherry/mRFP1 and mScarlet in our three- and four-colour reporter, we used a 580 nm laser line as excitation wavelength (Em. 590-650nm). This favoured the excitation of mCherry and mRFP1 (94%; 97%) over mScarlet (72%) (https://www.fpbase.org/compare/mrfp1,mcherry,mscarlet/) and allowed to efficiently co-visualize and discriminate the three corresponding fluorescent reporters emitting in the same spectral channel (Fig. S7). As an alternative, the red FP mKate2 could be used instead of mScarlet, as its spectral and photophysical properties are closer to those of mCherry (https://www.fpbase.org/compare/mcherry,mkate2/, Starling et al. 2023). Its lifetime of 2.5 ns, though, would reduce the dynamic range available to detect and quantify differences in reporter abundance. mRED7 (τ = 0.7 ns) and tagRFP-T (τ = 2.3 ns) have also been characterized as suitable FPs to use in combination with mScarlet and mCherry to label up to three spatially separated compartments in transiently transformed protonema cells of *Physcomitrium patens* (Aoyama et al. 2023). Although more complex Phasor-based separation of three, or even four, overlapping fluorescent species within a single spectral channel has been reported in non-plant systems (Vallmitjana et al. 2020; Frei et al. 2022b; Rahim et al. 2022), the efficient isolation of spatially overlapping fluorescent species is considered more challenging (Gonzalez Pisfil and Dietzel 2025).

The Phasor position of a fluorophore acts as a molecular fingerprint and allows to identify, separate, and quantify different molecular species. A fluorophore’s lifetime, though, is influenced by its molecular surroundings. By calculating the mean G and S coordinates of FPs targeted to different cell compartments, we demonstrate that both green- and red-emitting FPs show specific Phasor signatures depending on their subcellular localization, with a prominent shift towards shorter and multiexponential decays in the chloroplasts (Fig. 6 and Fig. 7). Such shift is sufficient to discriminate the same FP when localized to either the chloroplasts or the nucleus (Fig. 8A-C and S13). Consistent with our findings, differences in fluorescence lifetimes exhibited by chemically identical fluorophores targeted to different subcellular localizations were recently employed to achieve multiplexing using synthetic fluorescent probes (Frei et al. 2022a). Our data not only exemplify a workflow to precisely determine and compare the Phasor position of FPs, or other fluorescent species, in different molecular environments, but also emphasize the importance of taking into account the influence of those microenvironments on a fluorophore’s lifetime. While several probes have been specifically designed to take advantage of such sensitivity for measuring viscosity, pH or temperature in different biological systems (Okabe et al. 2012; Michels et al. 2020; Rennick et al. 2022), lifetime shifts can represent a confounding effect, especially when conducting quantitative studies using FPs. For instance, to reliably estimate the interaction between two molecular targets in FLIM-FRET studies, the donor’s lifetime in the presence and absence of the acceptor should be measured in a similar molecular environment, or at least in the same cellular compartment. If the donor changes localization upon interaction with the acceptor, it is advisable to measure the donor-only lifetime in the presence of an untagged version of the acceptor target.

Label-free approaches measuring the lifetime of endogenous fluorescent species are increasingly applied as they offer the possibility to visualize cells and tissues without the need of dyes, stains, or fluorescent labels. By imaging the endogenous fluorescence of lignin in fresh tomato root cross-sections, we show that FLIM enables to distinguish lignin depositions occurring in different tissues and exemplify how the Phasor approach can be used to characterise their peculiar decay (Fig 9-10). The different lifetime signatures we identified may be explained by a different composition and compaction of the polymer, and/or by the different lignification patterns. Lignin is indeed a complex aromatic polymer mainly consisting of p-Hydroxyphenyl (H), Guaiacyl (G) and Syringyl (S) units and its specific spatial patterning and monomeric composition confer peculiar properties to different cell types (Barros et al. 2015; Dixon and Barros 2019). While lignin is deposited within the matrix of cellulose and hemicellulose in secondary cell walls of the xylem, providing mechanical strength, rigidity and hydrophobicity, Casparian strips are rather dense, local lignin impregnations of the primary cell wall extending across the middle lamella acting as a waterproof apoplastic barrier (Emonet and Hay 2022). Interestingly, a previous study comparing the fluorescence lifetime of model lignin compounds containing almost pure lignin to artificial lignin–carbohydrate composites showed that increasing carbohydrate content leads to an increase of lignin fluorescence lifetime from 0.5-1 ns to 3 ns. By imaging the autofluorescence of latewood tracheids in poplar with FLIM, the authors further suggest that the lignin components that have short and long fluorescence lifetimes may correspond to “dense” (or concentrated) and “loosely” packed lignins, respectively (Zeng et al. 2015). This fits well with the short and long lifetimes we measured in the Casparian strips and xylem of tomato, respectively (Fig. 9-10). We observed longer fluorescence decays in metaxylem compared to protoxylem (Fig. 10D). This may reflect an increased S/G lignin ratio, as recent work has linked higher syringyl content with longer fluorescence lifetimes during protoxylem differentiation in Arabidopsis roots (Escamez et al. 202 1). On the other hand, it may be correlated to the different lignification patterns (reticulate or spiraled, respectively) characterizing these two tissues (Emonet and Hay 2022). Lignin composition and structure, however, can differ not only between cell-types but also in between plant species (Whetten et al. 1998). A more precise label-free biological description of autofluorescent structures may require the use of additional techniques. This has been exemplified when combining FLIM with Raman microscopy, where characteristic lifetime shifts accompanied changes in lignin chemistry, such as the replacement of alcohol groups with corresponding aldehydes in the polymer (Wightman et al. 2019). Such approaches in combination with using different treatments and/or plant mutants will foster our understanding of the biological nature and functions of these depositions.

**Figure 10.**
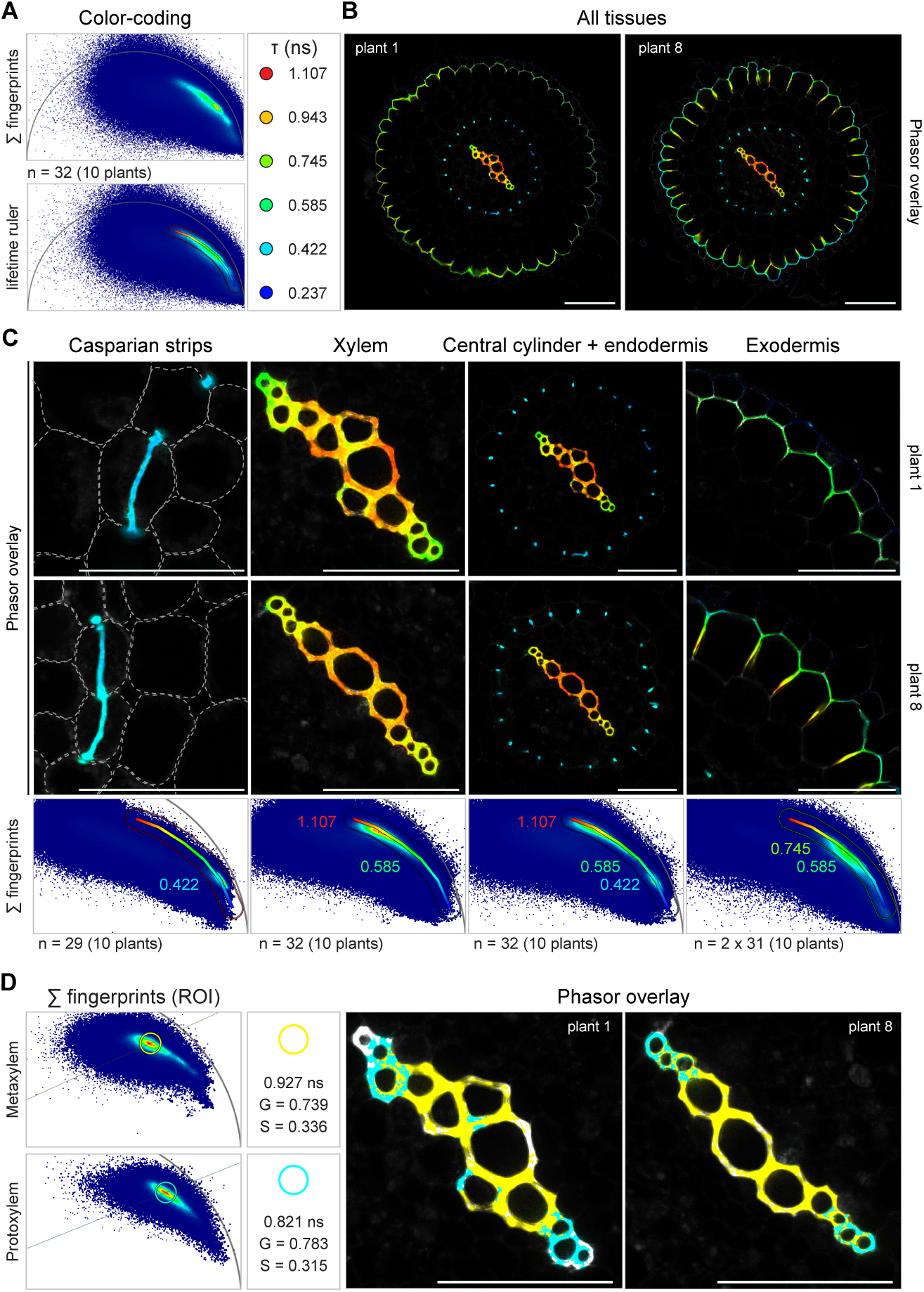
Phasor fingerprinting of the endogenous fluorescence. **A)** An all-encompassing colour-code was defined by positioning a lifetime ruler along the Phasor distribution resulting from the overlay of the Phasor fingerprints (Σ fingerprints) from all the images of entire cross-sections from the sample series. Lifetime values (τ, ns) of the outer and intermediate positions defined by the ruler are reported. **B**) Comparison of the endogenous fluorescence in two cross-sections from different plants by applying the Phasor colour-code to single confocal FLIM images. The Casparian strips (cyan) and the metaxylem (orange/red) exhibited the shortest and the longest decays, respectively, while the exodermal PLC (outer ring) and the protoxylem (terminal two-three cell files of the xylem associated to rather intermediate lifetimes (yellow/green). **C**) Phasor analysis and representative confocal images showing the tissue-specific Phasor fingerprints (Σ fingerprints) of lignin depositions in isolated regions of the cross-sections. Lifetime values defining the Phasor distributions and the corresponding colour-coding range on the Phasor overlay are reported on each fingerprint. The dashed white lines marking the cell borders in the Casparian strip images were drawn based on the corresponding bright-field images. **D**) Phasor positions of meta- (yellow) and proto- (cyan) xylem defined by centering the cursors on the Phasor distributions obtained by overlaying the fingerprints of regions of interest (ROIs) selected on the corresponding tissue (four central and two terminal cell files, respectively). Images are colour-coded according to the Phasor cursors. Their G and S coordinates together with the lifetime values (ns) are provided. n = number of images. Data are from two independent replicates. Scale bar = 100 μm (all tissues), 50 μm (Casparian strips, xylem, central cylinder + endodermis, exodermis).

In summary, our data illustrate how FLIM combined with Phasor analysis can be used for live-cell multiplexing and the discrimination of endogenous fluorescence in plants. The approach presented here was established using a TCS SP8 WLL Leica FALCON system, with Phasor analysis performed using an offline version of the Leica LAS X Control Software (version 4.8.2). This variant, which is integrated into the latest-generation “STELLARIS” FLIM platforms, enables the computation of Phasor cloud centres and the extraction of the corresponding G and S coordinates, which can be exported as Excel files and plotted to quantitatively support the analysis. Comparable functionality is available in the open-source software PhasorPy (https://www.Phasorpy.org/docs/stable/), which has been developed to facilitate Phasor-based image analysis across diverse microscopy platforms (Malacrida 2023). Although some of the features used in this study are specific to the Leica software (e.g., the “separation” function within the Phasor workflow), systematically reporting Phasor cloud coordinates—parameters that uniquely and unambiguously define fluorescent species—may help reduce compatibility constraints and support broader accessibility and reproducibility of FLIM data within the scientific community.

## MATERIALS AND METHODS

### Plant material and growth conditions

*N. benthamiana* seeds (∼30) were sown in 6 × 6 cm square pots filled with a soil–vermiculite mixture and transferred to a 22 °C growth chamber under a 16 h/8 h light/dark photoperiod. After one week, seedlings were transferred in pairs to new pots containing the same substrate and grown for an additional 4–5 weeks before being used for agroinfiltration. During this period, plants were watered three times per week. Seeds of *M. truncatula* wild-type (ecotype Jemalong A17) were soaked in sulfuric acid (H_2_SO_4_) 96% for 8 min, rinsed six times with sterile tap water and then surface sterilized with a solution of 1.2% sodium hypochlorite (NaClO) and 0.1% sodium dodecyl sulfate (SDS) for 1 min. Following six rinses with sterile tap water, seeds were transferred to 1% agar/water plates and vernalized at 4 °C in dark for 4–5 days. Seeds were then moved to 24 °C in the dark for 10 h to induce germination before being used for hairy root transformation. *S. lycopersicum* (cv. Moneymaker) seeds were surface sterilized with 70% ethanol for 3 min and then treated with a 6% (v/v) sodium hypochlorite (NaClO) solution containing a 0.01% Tween 20 for 20 min at 20 rpm on a rotating wheel. Seeds were then washed 6 times with sterile tap water, the first time shortly to remove most of the sterilization solution and then each time for 15 min at 20 rpm on a rotating wheel to induce swelling. Seeds were then transferred to 12 cm x 12 cm squared plates (20 seeds x plate) containing 2.2 g/L Murashige and Skoog medium including vitamins (Duchefa) and supplemented with 0.5 g/L MES pH 5.8, 40 g/L sucrose and 0.9 g/L phytoagar (Duchefa). Plates were kept horizontally in a 24 °C growth chamber for 7–10 days with 16 h of light and 8 h of dark per day.

### Construct design

The constructs used in this study were assembled using the Golden Gate modular cloning system (Weber et al. 2011; Binder et al. 2014). Plasmids were designed and annotated using Benchling. Details of L1 and L2 constructs, including module composition and *Agrobacterium* strains, are provided in the Supplementary data, together with the annotated sequences of L2 constructs (GenBank format).

L1 constructs for expressing the histone genes of Medicago (*H3.1, Medtr8g092720/H3.3, Medtr4g097175*) were designed according to Batzenschlager et al. 2025 and assembled using the same L0 modules containing the corresponding promoters and open reading frames (ORF). To express the remorin genes from Arabidopsis (*REM1.3, AT2G45820*) and Medicago (*SYMREM1, Medtr8g097320*), the ORF cloned from a genomic DNA template or the coding sequence cloned from a cDNA template were used, respectively. The L1 were designed according to Konrad et al. 2014. To assemble the L1 for the expression of fluorescent reporters targeted to the ER, chloroplasts, mitochondria and peroxisomes, the signal peptides or targeting sequences reported in (Nelson et al. 2007) were used. The L1 construct for targeting mCitrine to the PM was assembled using the 2xPH-FAPP1 L0 module synthesized by Life Technology and employed in Lace et al. 2023. All L1 constructs generated using the Ampicillin-resistance vectors from (Weber et al. 2011) were further cloned into L2 plasmids containing a kanamycin resistance marker to allow selection in *Agrobacterium*. For the multicolour transcriptional reporter, the L0 modules containing the promoter sequences of *RPG* (Medtr1g090807) and *VPY* (Medtr6g027840) reported in Lace et al. 2023 were used.

### Transformation of *N. benthamiana* epidermal cells

Recombinant *Agrobacterium* strains carrying the plasmid of interest were grown in liquid LB media supplemented with appropriate antibiotics for 24 h at 28 °C. Bacterial cultures were pelleted by centrifugation (4000 rpm) for 5 min, resuspended in Agromix (10 mM MgCl_2_; 10 mM MES/KOH pH 5.6; 150 uM Acetosyringone) and mixed in selected combinations to reach a final OD_600_ of 0.05. To enhance the transient production of the desired proteins, an *Agrobacterium tumefaciens* strain (GV3101) carrying a plasmid for the production of the Tomato Bushy Stunt Virus (TBSV) silencing suppressor p19 (Jay et al. 2023) was added to each mix (final OD_600_ = 0.05). The bacterial suspension was incubated at room temperature (RT) in the dark for 2 h and then injected into the leaf apoplast by placing a 1 mL needleless syringe against the abaxial side of the second and third leaf of 4-5 weeks old *N. benthamiana* plants. Plants were grown under 16 h light/8 h dark at 23-24 °C for 48-72 h, after which 2-3 small disks were excised from the leaves using a biopsy puncher and vacuum-infiltrated with tap water prior to imaging.

### Hairy root transformation and rhizobial inoculation

Composite *M. truncatula* plants were obtained according to the procedure previously described in Boisson-Dernier et al. 2005. In brief, transgenic *A. rhizogenes* strain ARqua1 carrying the plasmid of interest was cultured in 5 ml liquid LB medium supplemented with the appropriate antibiotics for 24 h, until reaching an OD_600_ of approximately 0.5–0.7. Subsequently, 300 μl of the bacterial suspension was spread onto solid LB medium containing the appropriate antibiotics and incubated at 28 °C in the dark for 48 h. For transformation, the root meristem of germinated seedlings was excised with a scalpel, and the wounded region was gently dragged across the *A. rhizogenes* lawn before seedlings were transferred to plates containing solid Fahräeus medium supplemented with 0.5 mM NH_4_NO_3_. Seedlings were grown vertically in a controlled-environment chamber at 22 °C in the dark for 3 days, followed by 4 additional days under a 16 h/8 h light/dark photoperiod while keeping the root system shaded. Composite plants were subsequently transferred to fresh solid Fahräeus medium supplemented with 0.5 mM NH_4_NO_3_ and grown for 10 days at 24 °C under a 16 h/8 h light/dark photoperiod. Transformed roots expressing the fluorescent selection marker were identified using a stereomicroscope, and non-transformed roots were removed with a scalpel. Composite plants were then transferred to solid Fahräeus medium supplemented with 0.1 mM NH_4_NO_3_ and grown for additional 4 days at 24 °C under a 16 h/8 h light/dark photoperiod before proceeding with bacteria inoculation. The *Ensifer medicae* strain WSM419 (Domonkos et al. 2017) constitutively producing mRFP1 was kindly provided by Benjamin Gurion (LIPM, Université de Toulouse, Castanet-Tolosan, France). The rhizobial strain was cultured in 5 ml liquid TY medium supplemented with the appropriate antibiotics for 3 days at 28 °C. Subsequently, 100 μl of this culture was used to inoculate a fresh 5 ml TY culture, which was grown for an additional 24 h at 28 °C. Bacterial cells were harvested by centrifugation at 3000 rpm for 10 min, washed once with tap water, and resuspended in tap water to a final OD_600_ of 0.01. Composite plants grown on solid Fahräeus medium supplemented with 0.1 mM NH_4_NO_3_ were then inoculated by applying 200 μl of bacterial suspension to each root system. Prior to inoculation, the position of the apex of transformed roots was marked on the plates. At 4–6 dpi, root segments that had grown below the marked position were excised, mounted in water, and imaged using a confocal microscope to capture infection events.

### Root sections and staining

For tomato root sections, 1 cm root segments were excised from the area located at 2-8 cm above the root tip of 7–9-day-old plants, embedded in 7% agarose and sectioned using a Vibratome (Leica VT1000 S) to obtain 70 µM-thick sections. Sections were then mounted in water for imaging of the endogenous fluorescence with FLIM. The lignin and cellulose dual staining was performed on a subset of root sections after FLIM imaging, by incubating them directly on the slide with 100 μl of a 0.05% (w/v) basic fuchsin solution (Sigma Cat# 857343-100g) for 10 min, and subsequently with 100 μl of a 0.01% (w/v) calcofluor white (Fluorescent Brightener 28, MP Biomedicals) solution for 5 min, with each staining step being followed by 3 washing steps of 5 min.

### Confocal Laser-Scanning Microscopy (CLSM) and Fluorescence Lifetime Imaging Microscopy (FLIM)

Confocal fluorescence microscopy and FLIM were performed on a Leica SP8 FALCON microscope equipped with a TCS SP8 X scanhead, a SuperK White Light Laser (WLL), a GaAsP-hybrid detector (HyD SMD), and a 20 x/0.75 water immersion lens (Leica Microsystems, Mannheim, Germany). The following spectral windows were used to acquire the fluorescent signal from fluorophores: 488 nm excitation / 500-550 nm emission (EGFP and NowGFP), 514 nm excitation / 520-570 nm emission (mCitrine and Venus), 561 nm excitation / 580-630 nm emission (mCherry and mScarlet in *N. benthamiana* epidermal cells) 580 nm excitation / 590-650 nm emission (mCherry, mScarlet and mRFP1 in *M. truncatula* transgenic roots). The endogenous fluorescence of root tissues was excited at 488 nm and the emission was collected at 500–550 nm. For the acquisition of basic fuchsin, the excitation was set at 571 nm and the emission collected at 575-615 nm. A 405 nm laser diode was used to excite calcofluor white, with the emission detected at 425-475 nm. To avoid crosstalk, different channels were acquired in sequential scanning between frames. Images of fluorescent reporters produced in *N. benthamiana* epidermal cells were acquired with a pulse repetition rate of 40 MHz and an accumulation of 1000 photons per pixel. The same repetition rate was used for imaging the three- and four-colour transcriptional reporter in *M. truncatula* transgenic roots colonized by fluorescent bacteria, but with an accumulation of 20 frames. To record the endogenous fluorescence of tomato root sections, a pulse repetition rate of 80 MHz was used, with an accumulation of 20 frames. All images are single focal planes, with the exception of the maximum intensity projections presented in Fig. 5B, Fig. S6B, Fig. S7A and the three-dimensional reconstructions presented in Fig. 9D.

### Image analysis and processing

All FLIM images were analysed using the LAS X FLIM Control Software version 4.8.2 (Leica Microsystems GmbH). The range of lifetimes used to colour-code FastFLIM images was adjusted to maximize contrast and is reported for every image. Separation of fluorescent species within single spectral channels was performed either via Phasor analysis or via global fitting of the intensity decay profile using n-exponential reconvolution. The number of components (n) used for curve fitting and the corresponding chi-squared (χ2) value is reported on the figures. A threshold of 30 photons was applied to generate the final images. For Phasor analysis, Phasor distributions (clouds or clusters) were computed by applying built-in wavelet-based denoising and first-harmonic Fourier transformation to fluorescence decay data, setting an intensity threshold of 6 photons per pixel. Phasor parameters (lifetime, G and S coordinates) of the different tools (cursors, ratio tool, lifetime rulers) used to analyse the Phasor plots were extracted from the software as Excel tables and are reported in the panels. To quantitatively analyse the Phasor signature of green- and red-emitting FPs in different subcellular compartments, either a single ROI (nucleus) or 8 to 10 ROIs (ER, cytoplasm, PM, mitochondria, peroxisomes, chloroplasts) were selected on each image (shown on the representative images presented in the figures). Cloud centres were computed with the built-in function of the software, the corresponding G and S coordinates were extracted as Excel tables and then plotted on a graph using GraphPad Prism software (GraphPad Software Inc), together with the average calculated in Excel. The number of images/cells used for the analysis is reported in the figures. All data obtained from both curve fitting and Phasor analysis presented in the article are provided in the Supplementary data. Intensity-only images and component images obtained from species separation using the Phasor approach or multiexponential curve fitting were further processed in Fiji (ImageJ) (Schindelin et al. 2012) for look-up table (LUT) adjustment, brightness and contrast optimization, scale bar annotation and generation of maximum intensity projections. For this purpose, datasets were saved after FLIM analysis as separate image series in Leica .lif format. When opened in Fiji, these series provide both the intensity-only image (8-bit) and component-specific intensity images (32-bit), representing the fractional contribution of each lifetime population. Both can be processed as standard confocal images. The adjusted minimum and maximum intensity values of the images corresponding to the isolated components are reported. For root cross-sections, visualization of 8-bit intensity data of the endogenous fluorescence was performed using the Magma LUT, ensuring perceptual uniformity and colourblind accessibility (https://bids.github.io/colourmap/). Standard confocal images of root cross-sections stained with basic fuchsin and calcofluor white were processed in Fiji (ImageJ). To analyse the decay of meta- and protoxylem, ROIs were selected on the four central or two terminal cell files, respectively. Phasor cursors were then centred on the two different Phasor distributions obtained by overlaying the Phasor fingerprints of ROIs from the two tissues.

### Statistical analysis

Statistical analysis and graphs generation were performed using GraphPad Prism software (GraphPad Software Inc). Data were subjected to a normality test to determine the statistical method to apply. Kruskal-Wallis followed by Dunn’s post-hoc test or unpaired t-test with Welch’s correction were applied as nonparametric or parametric tests, respectively. Raw data and results of the statistical analysis are provided in the Supplementary data.

## Acknowledgments

We would like to thank the entire team for constructive discussions and inputs on the project, and Casandra Hernández-Reyes, Nikolaj Abel, Guofeng Zhang for providing the EGFP-SYMREM1, the mCherry-REM1.3 and the mCherry-ER constructs, respectively. We would also like to thank Eija Schulze, Soraya Wilke Saliba and Rosula Hinnenberg for their continued technical help, Magdalini Tsitsikli for providing valuable feedback during figure preparation and Henriette Rübsam for the co-supervision of a Master student involved in the project. We are also grateful to Leica for generously providing access to the LAS X FLIM Control Software version 4.8.2. We also acknowledge the staff of the Life Imaging Center (LIC) at the Hilde Mangold House (HMH), Albert-Ludwigs-University of Freiburg, for their support with confocal microscopy. This work was conducted within the Collaborative Research Center 1381 (Sonderforschungsbereich SFB1381) under the project ID 403222702 and project 414136422 (T.O.) both funded by the German Research Foundation (DFG).

## Author contributions

Experimental design (B.L., T.O.), data collection (B.L., P.K., M.B., V.F., D.M.), analysis (B.L., P.K., M.B., V.F., D.M.), supervision (L.R., T.O.), writing (B.L., T.O.)

## Supplementary Figures and legends

**Supplementary Figure S1.**
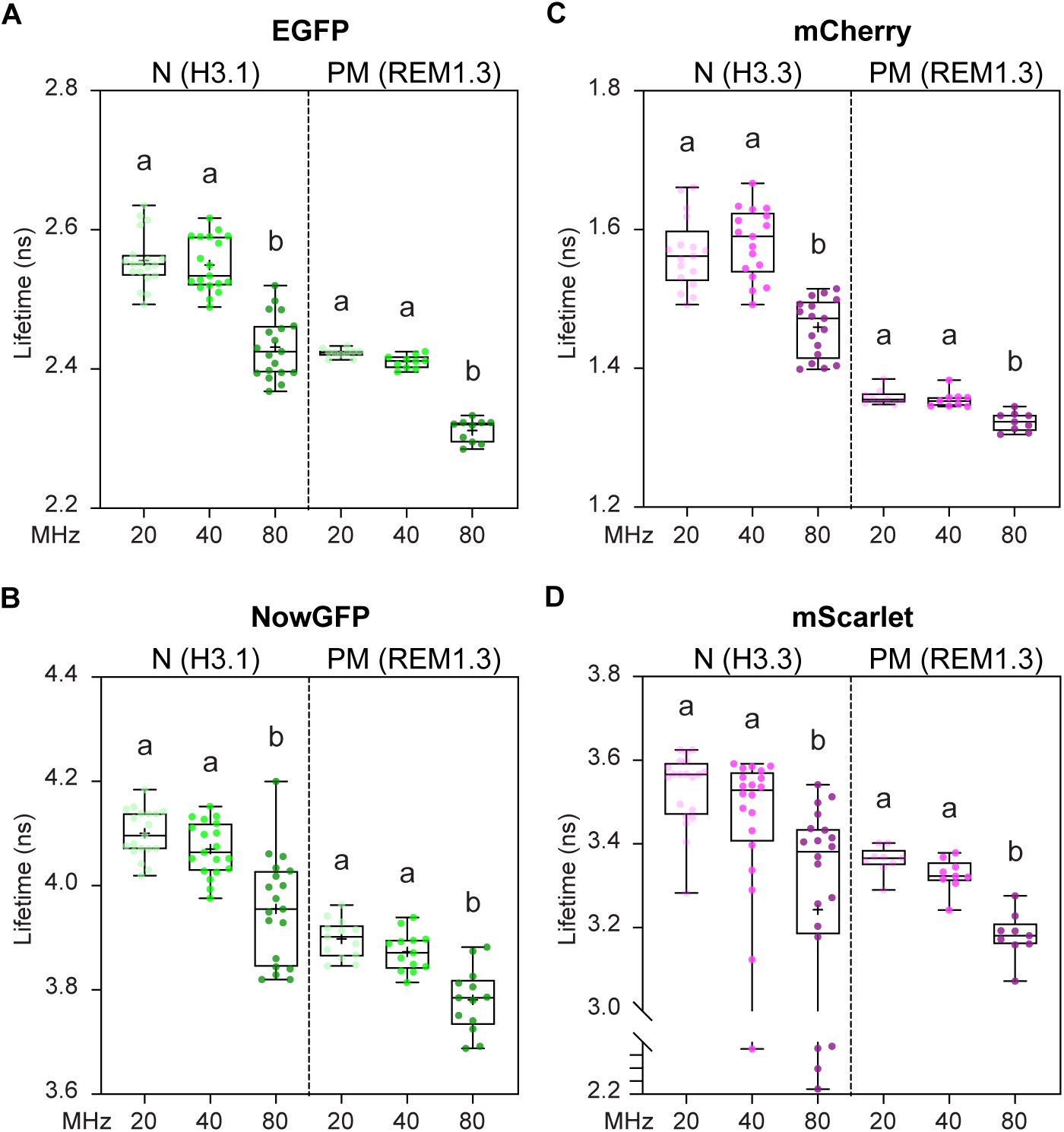
Lifetimes of FPs targeted to different subcellular compartments. **A-D)** Lifetime values (ns) of EGFP (**A**), NowGFP (**B**), mCherry (**C**) and mScarlet (**D**) targeted to the nucleus (N) or to the plasma membrane (PM) of *N. benthamiana* leaf epidermal cells. The same cell was imaged using 20, 40 and 80 MHz as repetition rate. Frames were accumulated until reaching 1000 photons per pixel and lifetimes were calculated by fitting a monoexponential model to the decay of either a single ROI on the nucleus or of at least 10 pooled ROIs on the PM. In the box plot, the top and bottom of each box represents the 75th and 25th percentiles, the middle horizontal bars indicate the median and the whiskers represent the range of minimum and maximum values. Crosses represent sample means. Letters indicate statistically significant differences according to Kruskal-Wallis multiple comparison analysis followed by a Dunn’s post-hoc test. n = 20 and 15 (**A**), 19 and 13 (**B**), 17 and 9 (**C**) 20 and 9 (**D**). Data are from two independent replicates.

**Supplementary Figure S2.**
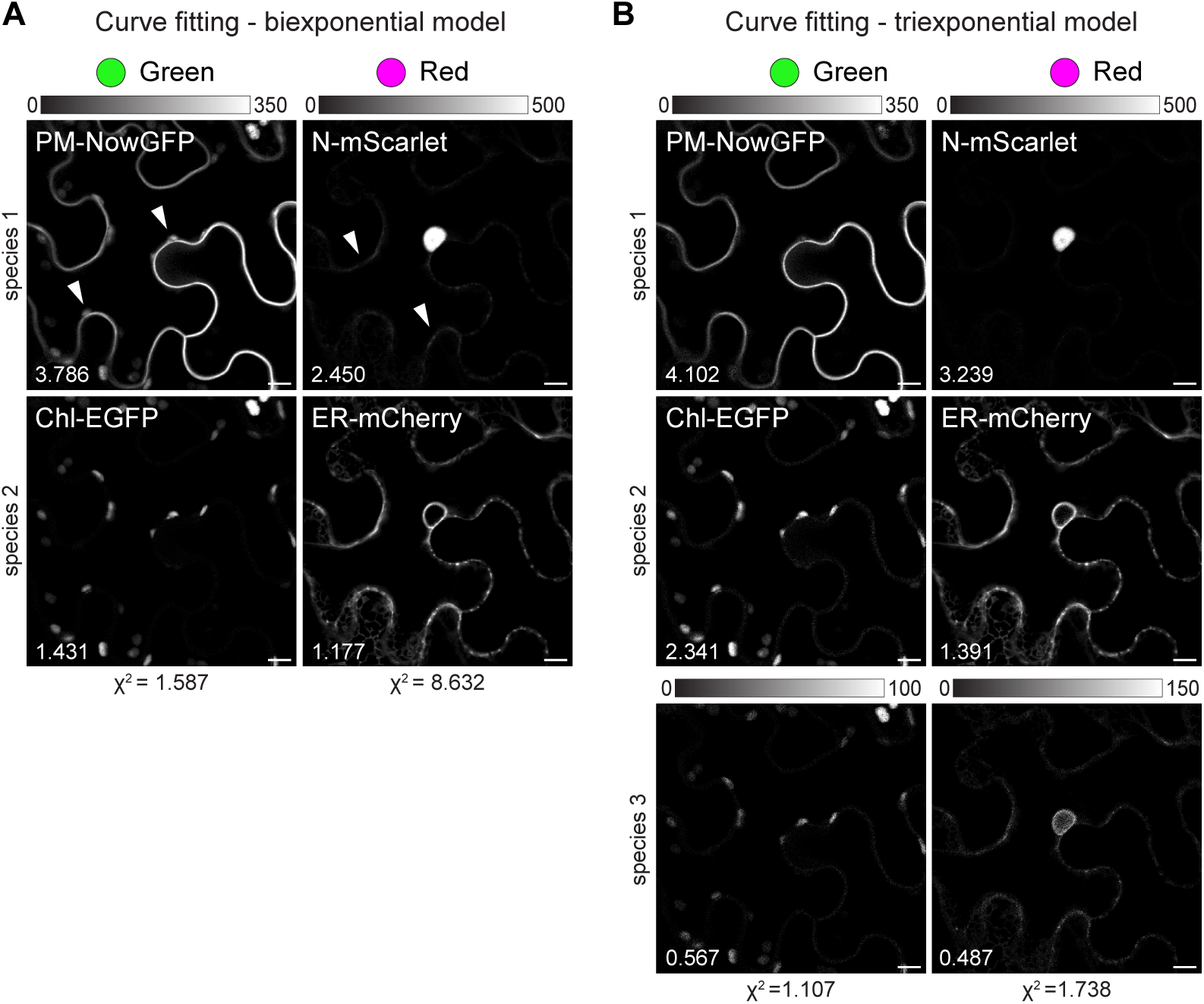
Separation of two pairs of fluorescent reporters in different spectral windows by curve fitting. **A-B)** Fluorescent species in the green and red channels were separated using a bi- (**A**) or tri- (**B**) exponential model to fit the decay. Images of the isolated species are shown in gray with the minimum and maximum intensity value indicated by the calibration bar. White arrowheads indicate pixels/subcellular structures assigned to the wrong species. chi square (χ2) and calculated mean lifetime values are provided. Original FLIM images are the same as those presented in Figure 2C. Scale bar = 10 μm.

**Supplementary Figure S3.**
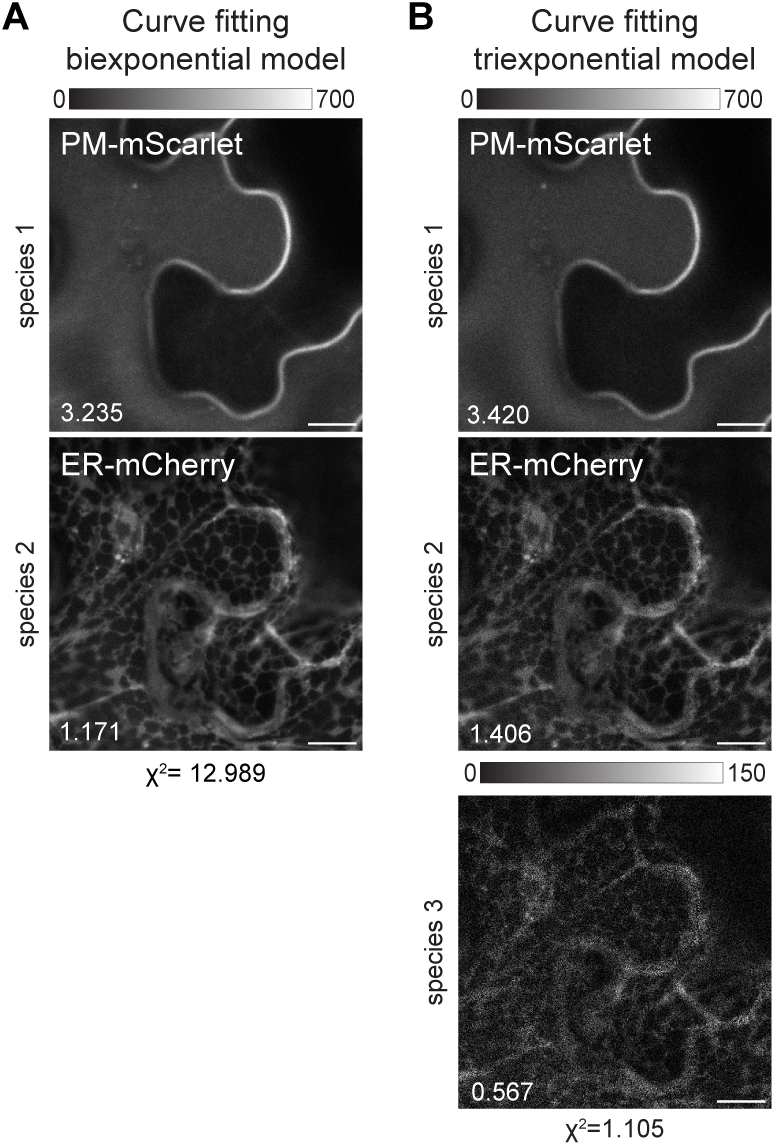
Separation of spatially overlapping fluorescent reporters by curve fitting. **A-B)** Fluorescent species were separated using a bi- (**A**) or tri- (**B**) exponential model to fit the decay. Images of the isolated species are shown in gray with the minimum and maximum intensity value indicated by the calibration bar. chi square (χ2) and calculated mean lifetime values are provided. Original FLIM images are the same as those presented in Figure 3C. Scale bar = 10 μm.

**Supplementary Figure S4.**
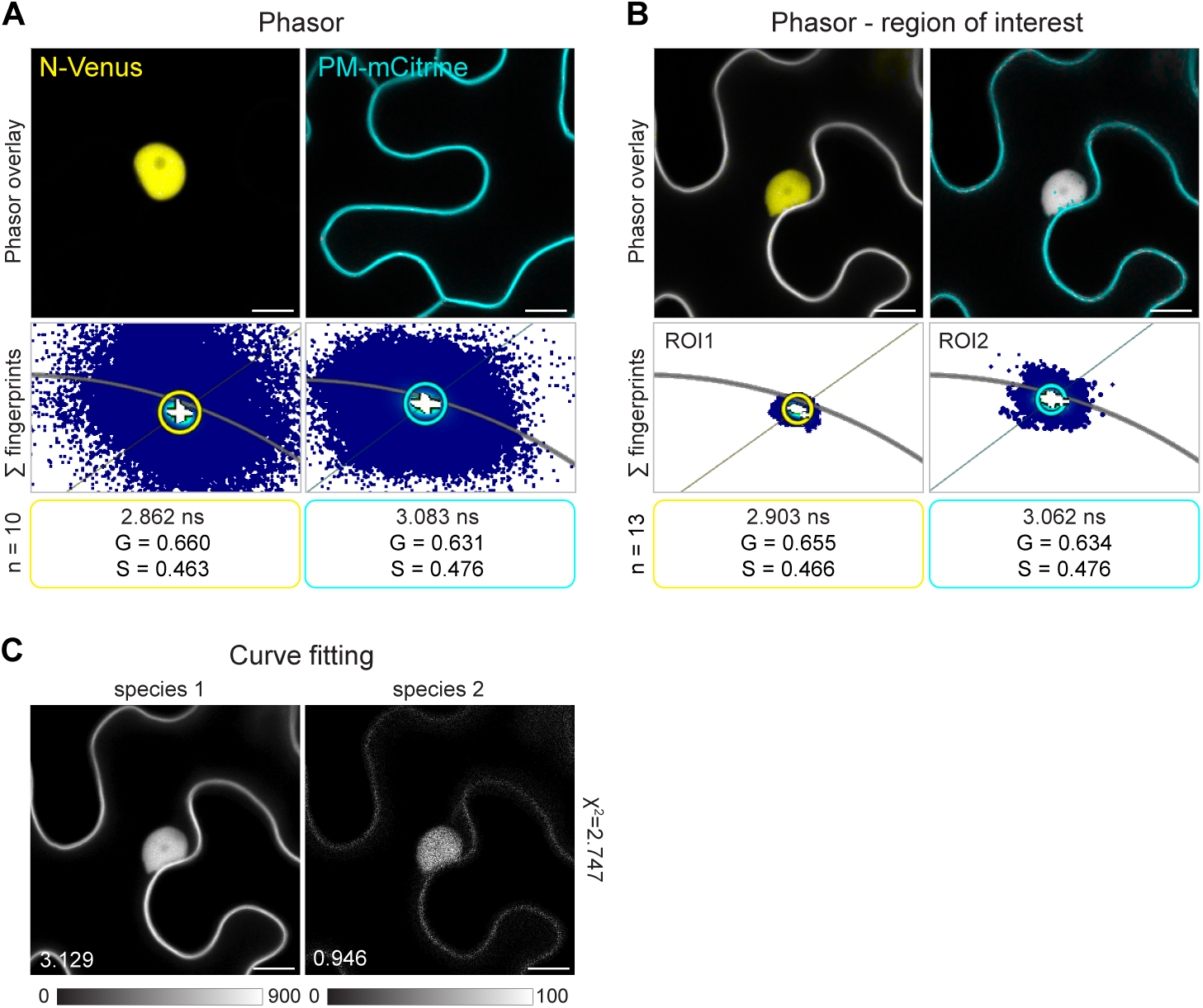
Characterization of yellow-emitting FPs with a small lifetime difference. **A-B)** Phasor analysis and representative confocal images of Venus (yellow) and mCitrine (cyan) targeted to the nucleus (N) and the plasma membrane (PM) of *N. benthamiana* leaf epidermal cells either separately (**A**) or simultaneously (**B**). The Phasor fingerprints of the images (**A**) or ROIs on the nucleus (ROI1) or the PM (ROI2) (**B**) were overlaid (Σ fingerprints) and the cursors were centered on the resulting clouds and used to colour-code the representative images. The G and S coordinates together with the lifetime values (ns) of the cursors are provided. n = number of images. Data are from at least two independent replicates. **C**) A biexponential model was used to fit the decay and separate the fluorescent species. Images are shown in gray with the minimum and maximum intensity value indicated by the calibration bar. chi square (χ2) and calculated mean lifetime values are provided. Original FLIM images are the same as those presented in Figure 3F. Scale bar = 10 μm.

**Supplementary Figure S5.**
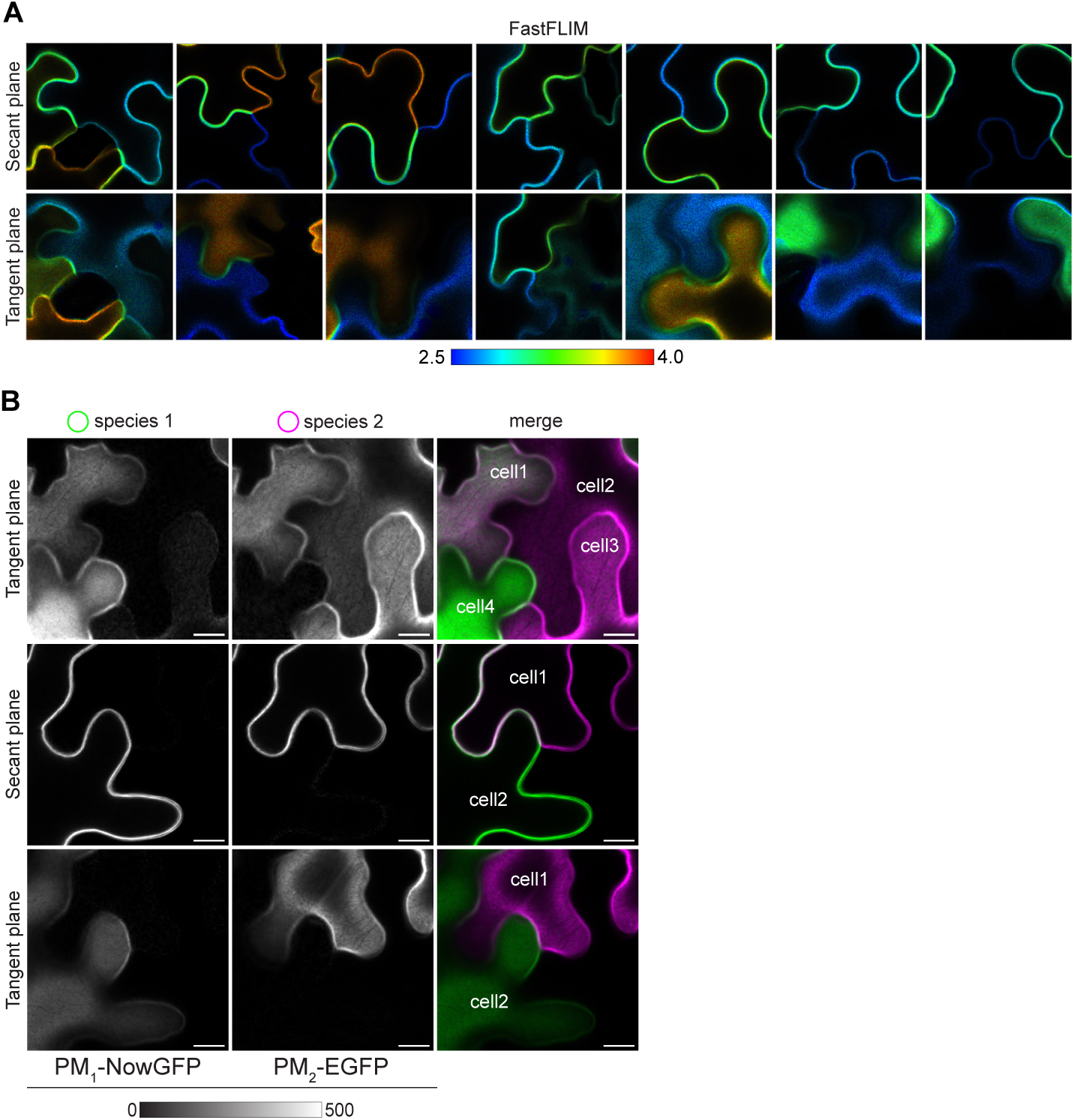
Lifetime-based separation of co-localizing fluorescent reporters. **A)** Additional set of FastFLIM confocal images of the boundaries between multiple epidermal cells acquired at the tangent and secant planes. **B**) Additional set of images showing the separation of the intensity contributions from the two PM-localized fluorescent reporters using the cursors defined in Figure 4I. Images of the isolated species are shown in gray with the minimum and maximum intensity value indicated by the calibration bar. Scale bar = 10 μm.

**Supplementary Figure S6.**
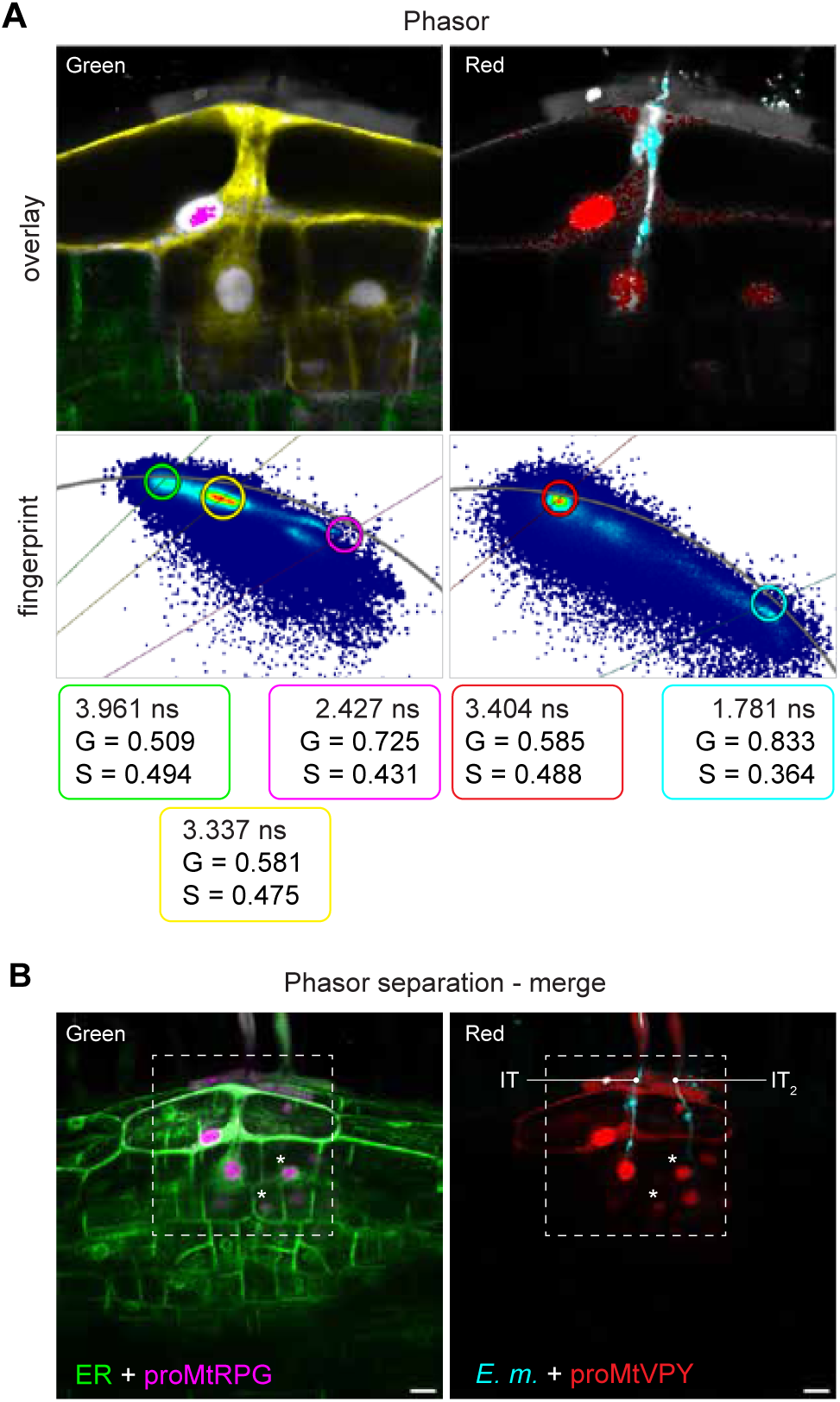
FLIM analysis of the three-colour transcriptional reporter during bacterial infection events in *M. truncatula* transgenic roots. **A)** Identification of the fluorescent reporters and mapping of their subcellular distribution based on the Phasor fingerprint. The yellow cursor was additionally placed on the Phasor distribution positioned on the mixing line between the NowGFP and the EGFP to map the overlap between the signal of the ER reporter and the cytoplasmic signal of the proRPG reporter in the image. The G and S coordinates together with the lifetime values (ns) of the cursors used to analyse the Phasor fingerprints are provided. **B)** FLIM images showing the same infection event shown in Fig. 5 imaged at two additional deeper focal planes. A second IT (IT_2_) close to the one previously analysed (IT) is penetrating the root tissues ahead of the cells where *RPG* and *VPY* appeared also co-expressed (asterisks). Images are maximum intensity projections of three focal planes obtained after Phasor separation of the individual species at each focal plane according to the Phasor cursors in (**A**). The merge of the species from each channel is shown, colour-coded according to the cursors. The white dashed box depicts the region imaged in (**A**) and in Fig. 5D. Scale bar = 20 μm.

**Figure S7.**
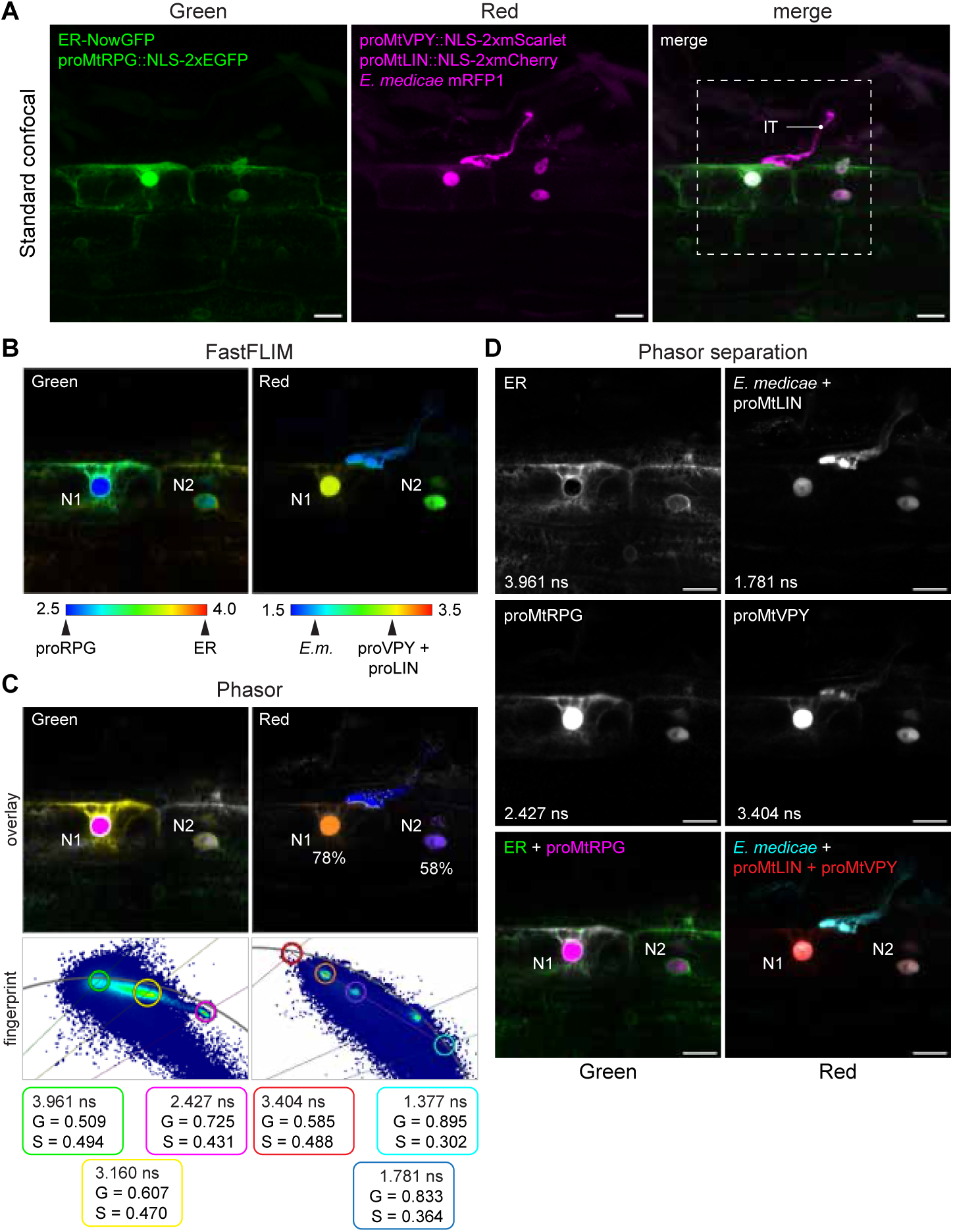
A four-colour transcriptional reporter to simultaneously monitor the co-expression of three different genes during bacterial infection of *M. truncatula* roots. **A)** Maximum intensity projections of a z-stack (14 steps, step size = 1 μm) showing an IT contacting a cortical cell in transgenic *M. truncatula* roots expressing the four-colour transcriptional reporter and inoculated with fluorescent bacteria. Images were acquired using standard confocal microscopy. The white dashed box in the merge depicts the area subsequently imaged using FLIM shown in (**B-D**). **B)** FastFLIM confocal images showing the co-activation of *RPG* (proRPG), *VPY* (proVPY) and *LIN* (proLIN) promoters in cortical cells underlying an IT. The five fluorescent reporters are mapped on the lifetime rainbow bar according to their expected lifetimes. *E.m*. = *Ensifer medicae* mRFP1. C) Identification of the fluorescent reporters and mapping of their subcellular distribution based on the Phasor fingerprint. The yellow cursor placed on the fingerprint of the green channel maps the overlap between the signal of the ER reporter and the cytoplasmic signal of the proRPG reporter in the image. The blue cursor indicates the signal from *Ensifer medicae* mRFP1. The orange and purple ratio tools in the red channel map the differential expression of VPY and LIN in the cell contacted by the IT (N1) and in the neighbouring one (N2). The G and S coordinates together with the lifetime values (ns) of the circular cursors used to analyse the Phasor fingerprints are provided. **D)** Separation of the intensity contribution from the fluorescent reporters in sub-images according to the Phasor cursors (green, magenta, red and cyan) in (**C**). Images of the isolated species are shown in gray. A minimum and a maximum intensity value of 0 and 800 or 400 were used to optimize the brightness and contrast of the green and the red channel, respectively. Merges are colour-coded according to the Phasor cursors (green, magenta, red and cyan) in (**C**). Lifetime values (ns) of the isolated species are annotated on the corresponding images. Images are representative of one independent replica with at least 5 infection events analysed. Scale bar = 20 μm.

**Supplementary Figure S8.**
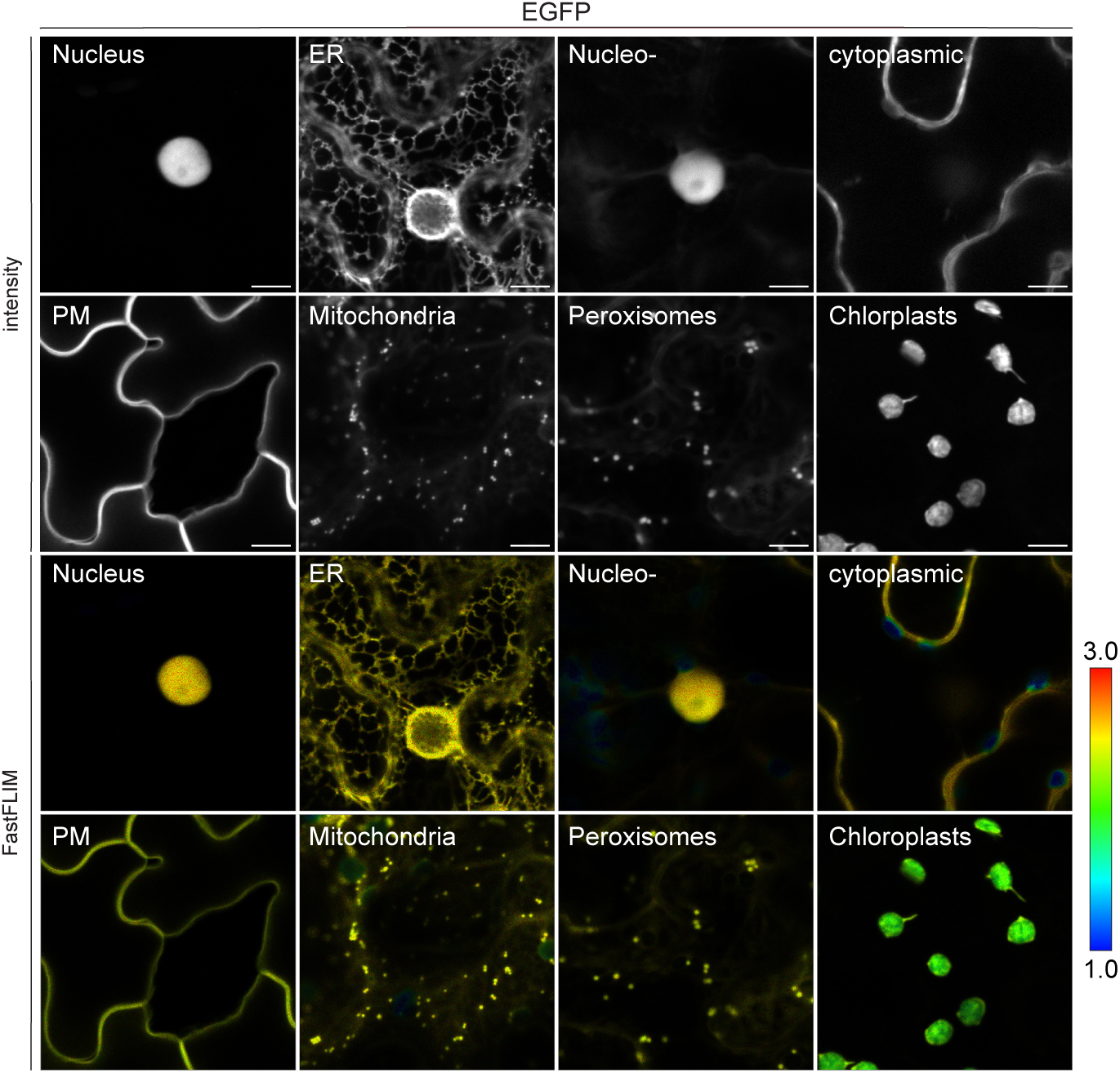
Intensity-only and FastFLIM confocal images of EGFP targeted to different subcellular compartments of *N. benthamiana* leaf epidermal cells showing the shorter lifetime exhibited by the FP in chloroplast compared to the other compartments. The average photon arrival time is colour-coded according to the rainbow bar. Scale bar = 10 μm.

**Supplementary Figure S9.**
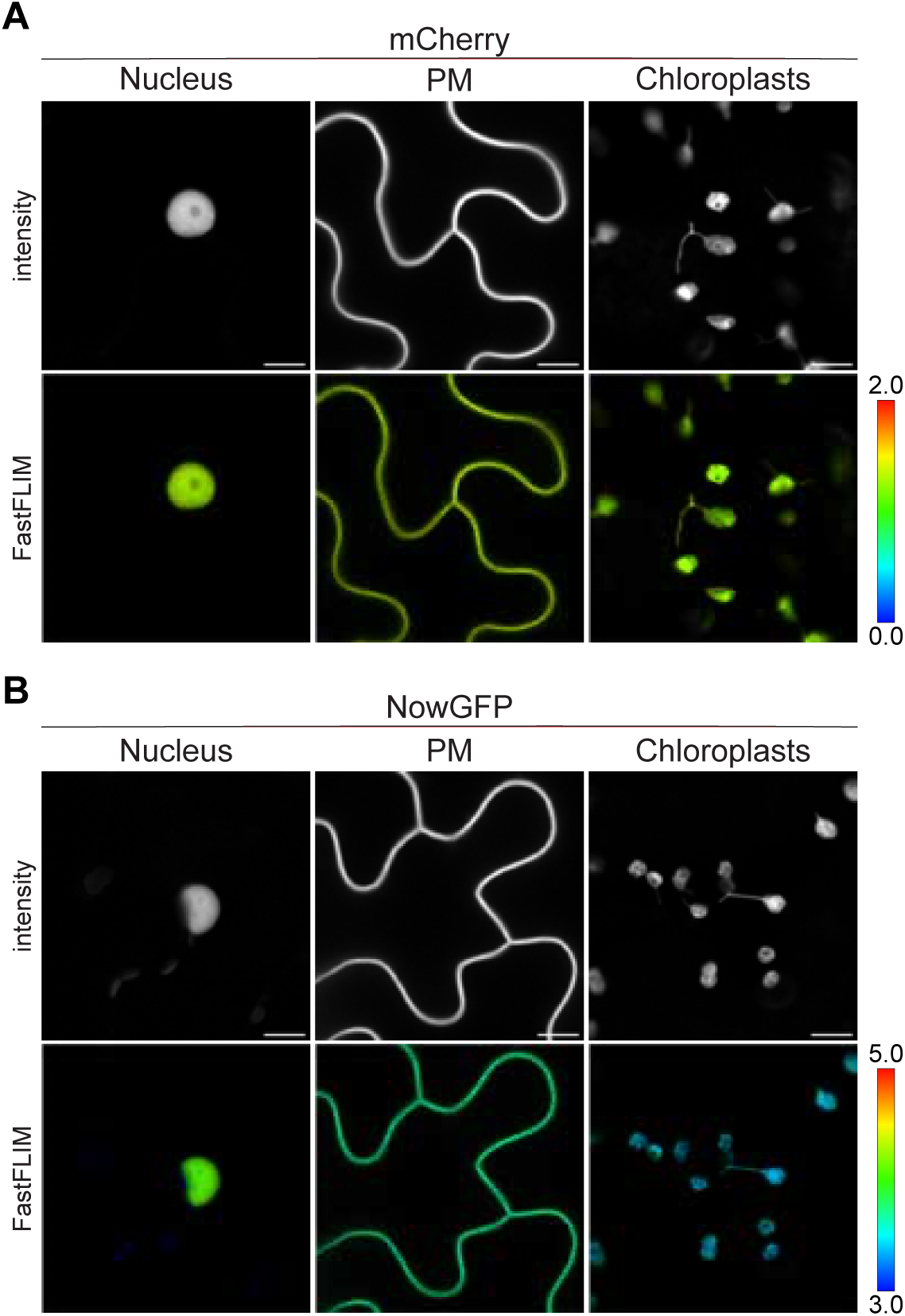
**A-B)** Intensity-only and FastFLIM confocal images of mCherry (**A**) and NowGFP (**B**) targeted to the nucleus, PM and chloroplasts of *N. benthamiana* leaf epidermal cells. The average photon arrival time is colour-coded according to the rainbow bar. Scale bar = 10 μm.

**Supplementary Figure S10.**
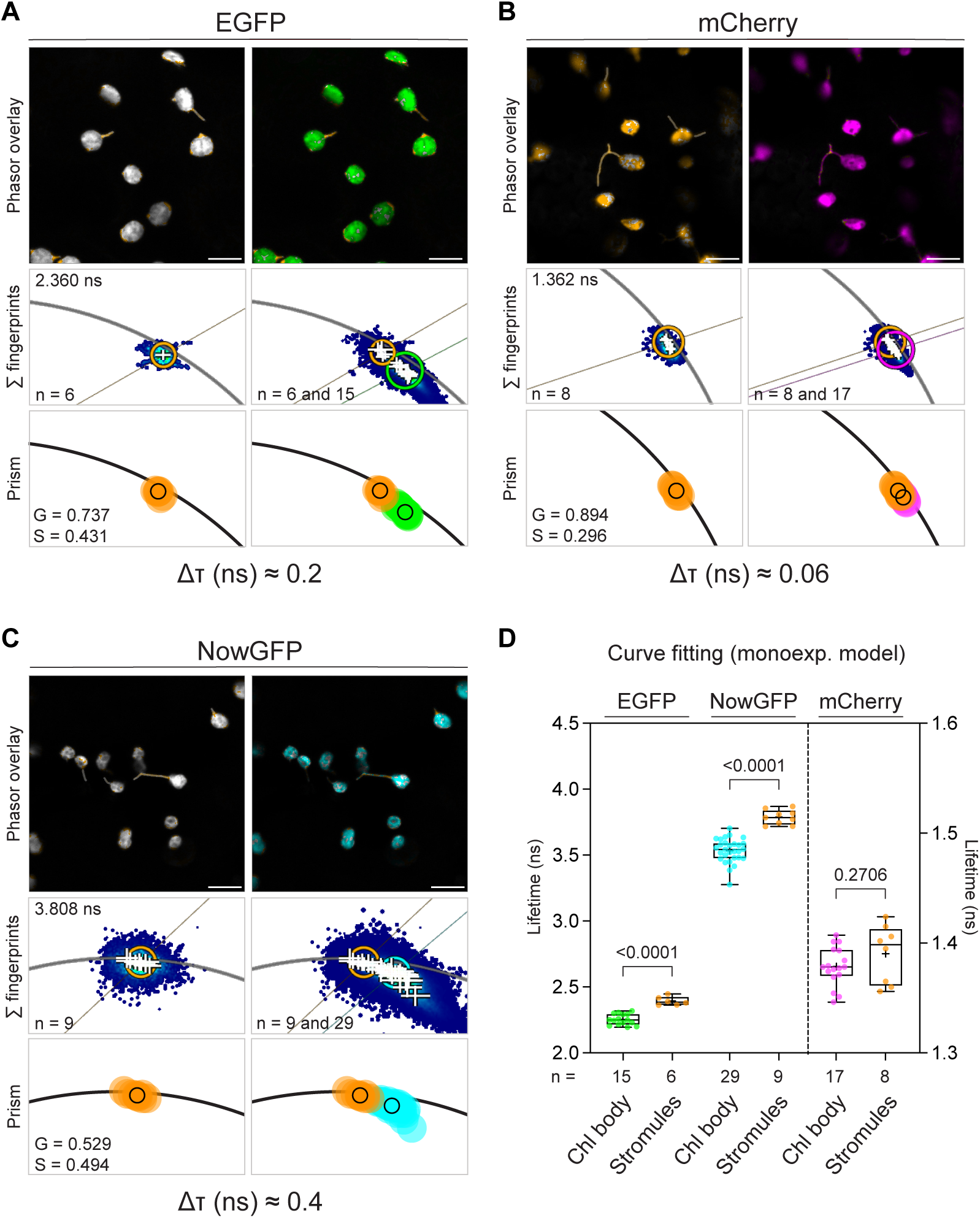
Specific decays associated to the stromules and body of chloroplasts. **A-C)** Phasor analysis and representative confocal images showing the distinct signatures of EGFP (**A**), mCherry (**B**) and NowGFP (**C**) in the stromules compared to the chloroplast bodies. The analysis was carried out as in Figure 5A (see legend), by additionally selecting ROIs on the stromules. The Phasor positions of the EGFP, mCherry and NowGFP in the stromules (orange circles) and in the chloroplast body (green, magenta and cyan circles, respectively) are shown together by overlaying their representative Phasor fingerprints (Σ fingerprints) and by plotting the average G and S on a graph. The lifetime values (ns) of the orange cursors used to analyse the Phasor fingerprint of stromules and the lifetime difference (Δτ) between the two sub-compartments are reported. Cursors and associated values of EGFP, mCherry and NowGFP in the chloroplast body are the same as in Figure 5A, 6A and 6B, respectively. **D**) Lifetime values (ns) of EGFP, NowGFP and mCherry in the chloroplast body (chl body) and the stromules calculated by fitting a monoexponential model to the decay of the corresponding ROI. In the box plot, the top and bottom of each box represents the 75th and 25th percentiles, the middle horizontal bars indicate the median and the whiskers represent the range of minimum and maximum values. Crosses represent sample means. An unpaired t-test with Welch’s correction was used to calculate statistically significant differences. n = number of images. Data are from at least two independent replicates. Scale bar = 10 μm.

**Supplementary Figure S11.**
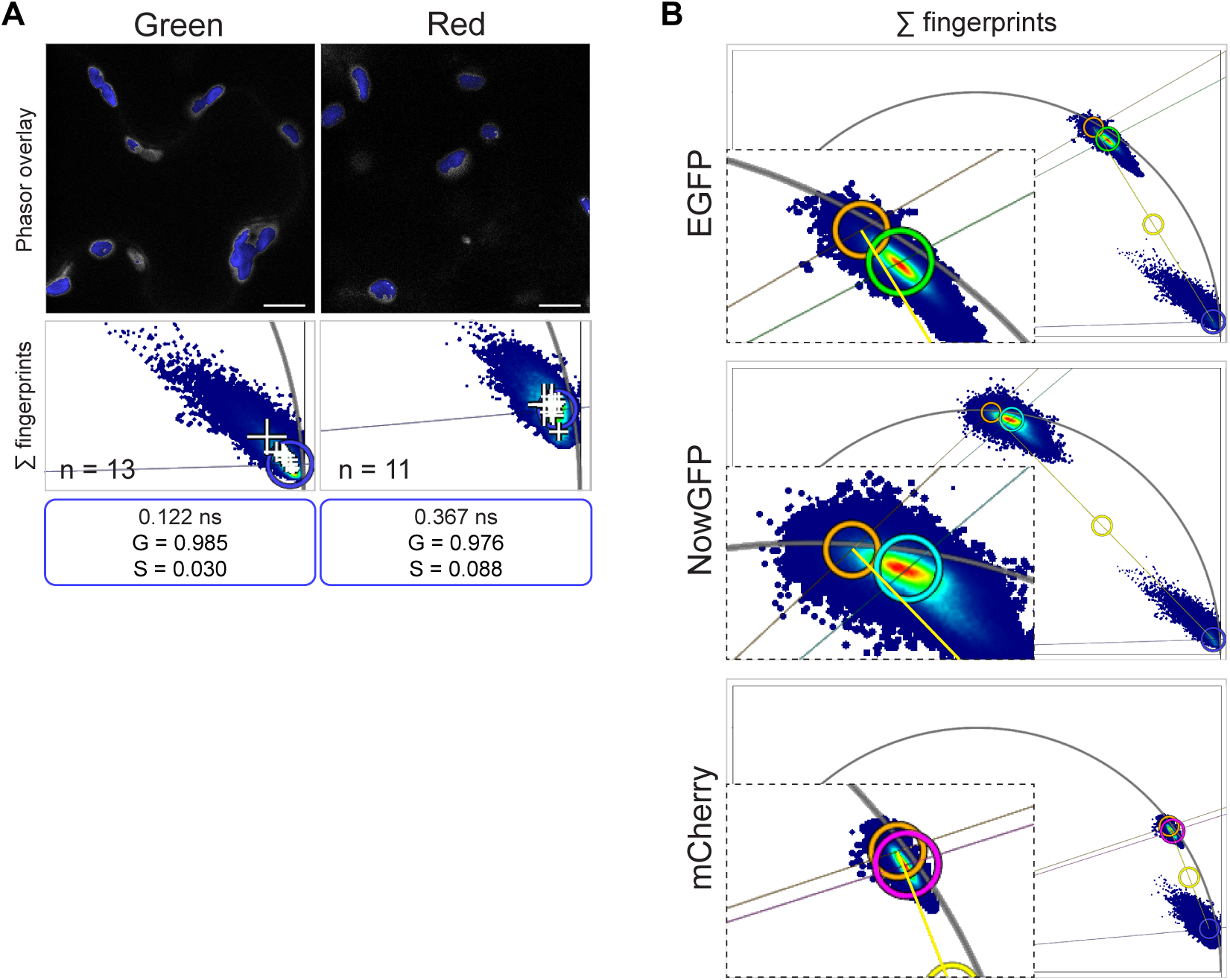
The chloroplasts-specific shift of FPs is not caused by the endogenous fluorescence. **A**) Phasor analysis and representative confocal images showing the Phasor position (blue cursor) of the endogenous fluorescence emitted by chloroplasts in untransformed *N. benthamiana* leaf epidermal cells. Images were acquired with the same excitation and emission parameters used for EGFP, NowGFP (Green) and mCherry (Red). **B**) The Phasor positions of both the EGFP (green cursor) and of the NowGFP (cyan cursor) in chloroplasts do not lie on the mixing line (yellow) connecting the positions of the stromules (orange cursor) and of the endogenous fluorescence (blue cursor). For mCherry, the two Phasor positions are too close for a reliable estimation. The G and S coordinates together with the lifetime values (ns) of the cursors used to analyse the Phasor fingerprint are provided. n = number of cells. Data are from at least two independent replicates. Scale bar = 10 μm

**Supplementary Figure S12.**
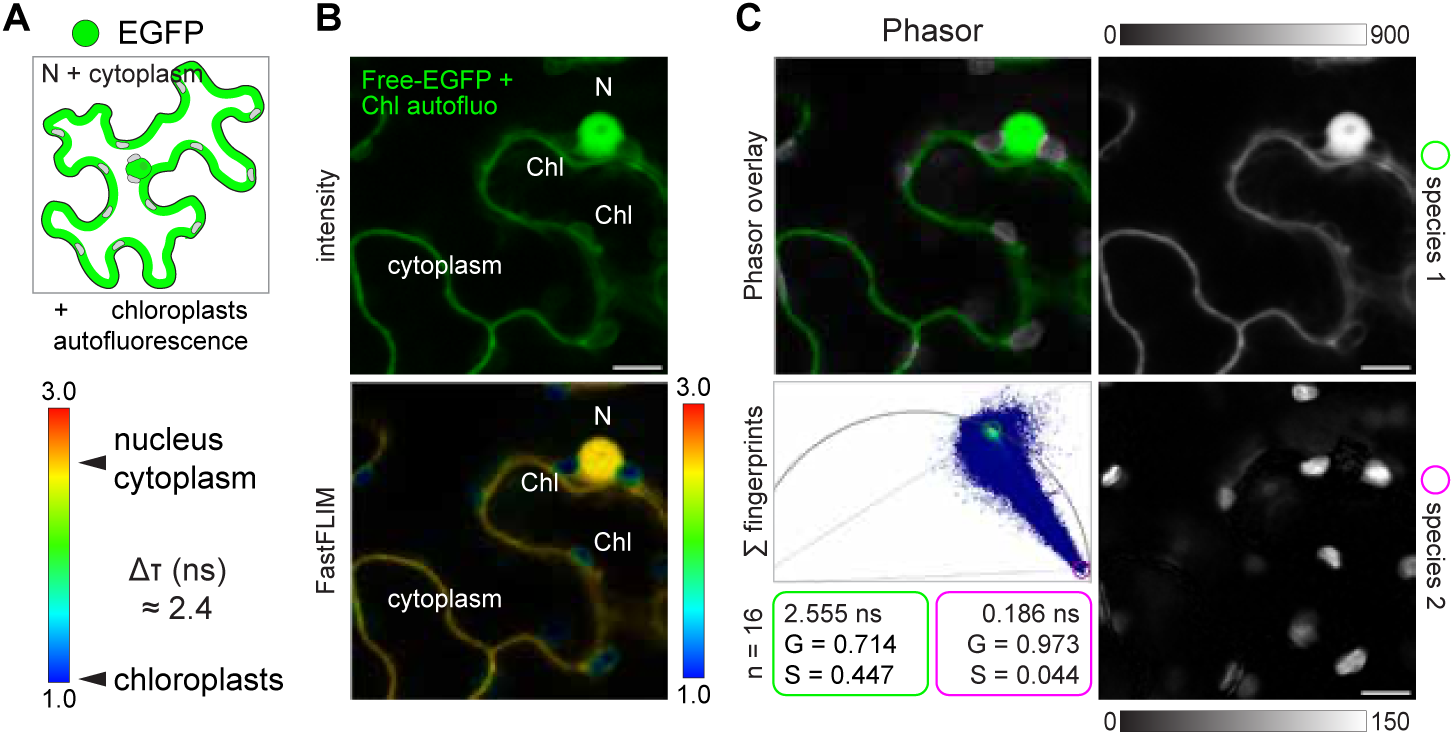
FLIM discriminates EGFP from chloroplast autofluorescence. A-C) Free-EGFP localized to the nucleus (N) and cytoplasm of *N. benthamiana* leaf epidermal cells (schematic in (**A**)) was used to exemplify the discrimination of chloroplast (Chl) autofluorescence based on the lifetime difference (Δτ). Chloroplasts are easily distinguishable in the FastFLIM image due to their short decay (**B**) and their signal can be isolated from the nucleo-cytoplasmic EGFP using Phasor analysis (**C**). Images of the isolated species in (**C**) are shown in gray with the minimum and maximum intensity value indicated by the calibration bar. The G and S coordinates together with the lifetime values (ns) of the cursors used to analyse the Phasor fingerprint are provided. n = number of cells. Data are from two independent replicates. Scale bar = 10 μm

**Supplementary Figure S13.**
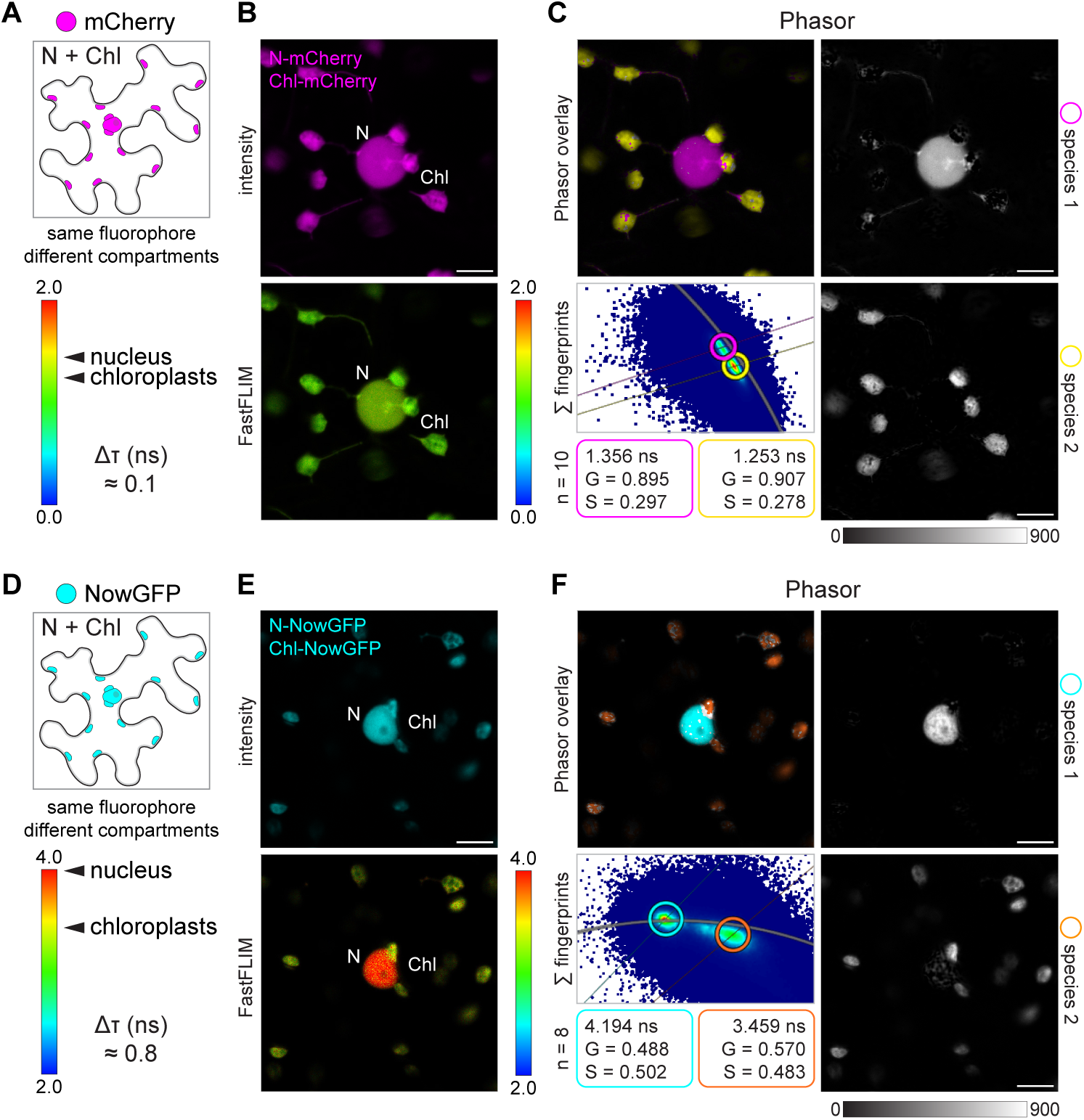
Phasor separation of the same FP targeted to different subcellular compartments. **A-B)** mCherry (**A**) and NowGFP (**B**) were simultaneously targeted to the nucleus (N) and the chloroplasts (Chl) of *N. benthamiana* leaf epidermal cells to test if the lifetime shift (Δτ) supports discrimination of the two compartments. **B-E)** Intensity-only and FastFLIM confocal images of an epidermal cell showing the discrimination of the same FP based on lifetime values (rainbow bar). **C-F)** Phasor separation of the chloroplasts- and nuclear-localized FPs mapping to distinct positions in the Phasor plot across cells (Σ fingerprints). Images of the isolated species are shown in gray with the minimum and maximum intensity value indicated by the calibration bar. The G and S coordinates together with the lifetime values (ns) of the cursors used to analyse the Phasor fingerprint are provided. n = number of cells. Data are from two independent replicates. Scale bar = 10 μm

